# Enhanced carotenoid photoprotection in Far-Red light acclimated *Chroococcidiopsis thermalis*

**DOI:** 10.64898/2025.12.23.696154

**Authors:** Elisabetta Liistro, Andrea Calcinoni, Alessandro Agostini, Samuel Pressi, Ester Marotta, Nicoletta La Rocca, Donatella Carbonera

## Abstract

Oxygenic photosynthesis is mainly driven by visible light in most photosynthetic organisms. However, some cyanobacteria strains can reversibly remodel their photosynthetic apparatus in order to rely exclusively on far-red photons. This acclimation, known as FaRLiP, requires the synthesis of red-shifted pigments, chlorophyll *d*, chlorophyll *f* and far-red allophycocyanin, that are incorporated in paralog subunits of the main photosynthetic complexes, namely photosystem II and photosystem I as well as phycobilisomes. In addition, some far-red–acclimating strains were also observed to show a rise in the carotenoid content and in the expression of genes involved in the biosynthesis of UV-shielding molecules, suggesting a counterintuitive photoprotective response concomitant with acclimation to lower energy wavelengths. In this work we investigated the response of *Chroococcidiopsis thermalis* to far-red light, and identified a robust set of photoprotective mechanisms associated with acclimation. Enhanced carotenoid/chlorophyll content ratios correlated with stronger carotenoid–chlorophyll triplet quenching, particularly pronounced for red-shifted chlorophylls. Non-photochemical quenching was higher and activated more rapidly than in cells grown under simulated solar light, and this was accompanied by enhanced antioxidant activity. Collectively, these findings indicate that the far-red acclimated cells, whose photosynthetic apparatus is not optimal under visible light, exhibit a strong photoprotected state, that we propose to be crucial under fluctuating irradiance conditions and during transitions from shaded to non-shaded environments.

**Highlights:**

- Far-red light photoacclimation (FaRLiP) in *Chroococcidiopsis thermalis* is associated with an increased carotenoid-to-chlorophyll ratio. The relative abundance of myxol-2’ fucoside, echinenone and β-carotene increased in cells acclimated to far-red light compared with cells acclimated to simulated solar irradiation.
- Upon establishment of FaRLiP an exceptionally strong chlorophyll triplet quenching by carotenoids was observed, and it was specifically involving red-shifted chlorophylls.
- When exposed to the same light treatment, non-photochemical quenching was higher in far-red-acclimated cells than in solar-acclimated. Moreover, following high visible-light stress, far-red-acclimated cells exhibited lower levels of reactive oxygen species. The FaRLiP photosynthetic apparatus is safeguarded by robust mechanisms of photoprotection.

## 1. Introduction

The majority of photosynthetic organisms rely on visible light for oxygenic photosynthesis, as far-red wavelengths are energetically insufficient to efficiently drive the process and only weakly absorbed by specific chlorophyll *a* molecules within Photosystem I (PSI) (Karapetyan et al., 2006; Morosinotto et al., 2003; Gobets & Van Grondelle, 2001). Nevertheless, in the last decades, several cyanobacterial species have been discovered capable of sustaining photosynthesis under monochromatic far-red light. Cyanobacteria, a diverse group of oxygenic photosynthetic prokaryotes, are found in a wide range of environments. Some species are able to thrive in subsurface environments – including stones, sediments, caves, biofilms, and symbiotic associations – where light availability is scarce, either in intensity or quality (Waditee-Sirisattha & Kageyama, 2022). These environments typically have limited visible light but are enriched in far-red light. The first cyanobacterium identified as capable of efficient oxygenic photosynthesis using far-red photons was *Acaryochloris marina*, a chlorophyll *d* bearing strain where this red-shifted pigment is expressed constitutively and accounts for up to 80% of the total lipid-soluble pigment content (Miyashita et al., 1996). In the following years cyanobacteria capable of synthesizing chlorophyll *f* (Chl *f*) besides chlorophyll *d* (Chl *d*) have been discovered (Antonaru et al., 2020, 2025; Gan et al., 2015). These species can remodel their photosynthetic apparatus to utilize far-red light through the so-called far-red light photoacclimation (FaRLiP) response, which is reversible depending on light conditions (Gan et al., 2014). FaRLiP-capable cyanobacteria are remarkably diverse in terms of ecology, geographical distribution and phylogeny, yet they colonize different ecological niches all characterized by far-red enriched spectra (Gan et al., 2015; Antonaru et al., 2020). Reported examples include hydrothermal species, e.g. *Leptolyngbya* sp. JSC-1 (Gan et al., 2014), endolithic species, e.g. *Chroococcidiopsis* sp. CCMEE 010 (Billi et al., 2022), intertidal species, e.g. *Synechococcus* sp. PCC7335 (Gan et al., 2015) and species isolated from soils, e.g. *Chlorogloeopsis fritschii* PCC6912 (Airs et al., 2014) and from stromatolites, e.g. *Halomicronema hongdechloris* (Chen et al., 2012). Interestingly, only two genera, *Kovacikia* (Zampieri et al., 2025) and *Leptodemis* (Shen et al., 2025) have so far been shown to include a high number of FaRLiP species.

The FaRLiP response is induced when cells are exposed to far-red enriched light spectra and it is enabled by a gene cluster of approximately 21 genes, with a certain diversity among species (Elias et al., 2024). The response involves the synthesis of new pigments, Chl *d*, Chl *f*, and far-red allophycocyanin (FR-APC), along with specific proteins that incorporate these pigments in a remodeled photosynthetic apparatus, including both photosystems and light harvesting antennae (Gisriel, Cardona, et al., 2022; Gisriel et al., 2023; Soulier et al., 2020; Trinugroho et al., 2020; M. Y. Ho et al., 2017; Gan et al., 2014).

During this acclimation, the photosystems expressed in white light are substituted with new photosystems made up of paralog FaRLiP subunits and incorporating also Chl *d* and Chl *f*, shifting the absorption and fluorescence emission of both photosystems far beyond 700 nm. A detailed overview of the structures of the photosystems expressed in far-red light can be found in Gisriel, 2024.

Briefly, in far-red light PSII (FR-PSII) the subunits D1, CP47, CP43, D2 and PsbH are FaRLiP paralogs. FR-PSII was found to harbor 30 Chl *a*, 4 Chl *f* and 1 Chl *d* and to have a long-wavelength electron donor (Nürnberg et al., 2018), which cryo-EM structures from *Synechococcus* sp. PCC7335 suggest may be represented by a Chl *d* in the Chl_D1_ position (Gisriel et al., 2022a). The FaRLiP paralog subunits in far-red light PSI (FR-PSI) are PsaA, PsaB, PsaF, PsaJ, PsaI and PsaL, coordinating at least 5 Chl *f*, located within the antenna (Gisriel et al., 2022b; Gisriel et al., 2021; Gisriel et al., 2020; Kato et al., 2020). A recent structural work from Consoli and collaborators (2025) suggested inclusion of one Chl *f* in the PSI reaction centre instead.

Interestingly, the FaRLiP trait has recently been demonstrated to have an ancestral origin in the evolution of the cyanobacteria phylum, emerging in the Paleoproterozoic, that was moreover preceded by the appearance and split of the *Acaryochloris* clade (Antonaru et al., 2025). Antonaru and colleagues sustain the relevance of photosynthesizing in shaded environments, enriched in far-red light, in the formation of microbial mats and stromatolites during the early Paleoproterozoic. The FaRLiP cluster has been then inherited vertically during the diversification of cyanobacteria thanks to its co-localization and the relatively simple regulatory system (Antonaru et al. 2025).

By enabling survival in niches sheltered from harmful UV and high-intensity visible radiation — at a time when the ozone layer was absent — FaRLiP may have represented a key ecological innovation in early cyanobacterial evolution.

Indeed, in addition to assembling a far-red harvesting photosynthetic apparatus, *C. fritschii* was reported to increase the production of mycosporine-like amino acids (MAAs) under far-red light, (Llewellyn et al., 2020). Typically, MAA biosynthesis is induced by ultraviolet radiation as a protective response against photodamage. MAAs absorb between 310 and 360 nm and function as both antioxidants and sunscreens (Geraldes & Pinto, 2021). Therefore, the FaRLiP response seems to be counterintuitively coupled to an increase in photoprotection. On the same line, a higher *in vivo* absorbance at blue–green wavelengths attributable to carotenoids has been observed in some FaRLiP strains acclimated to far-red light (Gan et al., 2015; Schmitt & Friedrich, 2024). *C. fritschii* was also found to have a higher carotenoid concentration when grown under white light combined with far-red light (Silkina et al., 2019). Carotenoids are known to participate in photoprotective mechanisms within photosynthetic systems too, and this specific role seems to be coupled to far-red light exposure, since their light harvesting properties are purposeless in the long-wavelengths regime. Despite these observations, the photoprotective activities accompanying FaRLiP have received relatively little attention. Photoprotection is fundamental for the survival of photosynthetic organisms, as excess light energy that exceeds the turnover of photosynthetic reactions can cause oxidative stress and damage to the photosynthetic apparatus, leading to either permanent photoinactivation or reversible photoinhibition (Derks et al., 2015; Melis, 1999). To cope with excessive light, oxygenic photosynthetic organisms display photoprotective strategies that are mainly operated by carotenoids. Carotenoids are crucial for non-photochemical quenching, NPQ, a series of mechanisms that dissipate the excessive energy (Arshad et al., 2022). In cyanobacteria, the fastest component of NPQ is operated by the orange carotenoid protein, OCP, that acts as an alternative energy sink downward phycobilisomes, PBSs, when reaction centers are closed (Wilson et al., 2006, 2012). OCP is a photoactivated protein carrying a carotenoid, typically 3-hydroxyechinenone, its precursors being echinenone or canthaxanthin (Niziński et al., 2025). When activated by absorption of blue-green light, OCP binds PBSs and the carotenoid collects the energy in excess, which is subsequently dissipated as heat (García-Oneto et al., 2024; Domínguez-Martín et al., 2022; Muzzopappa & Kirilovsky, 2020; Punginelli et al., 2009).

Under high light stress, carotenoids also directly quench triplet excited states of chlorophyll (^3^Chl), preventing their reaction with molecular oxygen and thereby inhibiting the formation of harmful ^1^O_2_ and other reactive oxygen species, ROS (Fiebig et al., 2023; Hashimoto et al., 2016; Polívka & Sundström, 2004). Carotenoids can also directly quench ^1^O_2_, and return to the ground state by releasing the energy via thermal dissipation (Hashimoto et al., 2016; Ramel et al., 2012).

To assess whether the FaRLiP response is paired with a peculiar photoprotective response, in this study we investigated the acclimation to far-red light in the FaRLiP strain *Chroococcidiopsis thermalis* PCC7203, with a particular focus on photoprotection. Using a combination of spectroscopic and biochemical approaches, we characterized the carotenoid content and evaluated the roles of these molecules in far-red acclimated cells. Photoprotection was assessed by detecting carotenoid triplet states – diagnostic of the pre-emptive quenching of chlorophyll triplet-derived oxidants – using ODMR spectroscopy. Complementary ROS measurements with a fluorescent probe revealed distinct photodamage susceptibilities in far-red versus solar light acclimated cells. Finally, analysis of NPQ activation provided further information on the photoprotective repertoire deployed by *C. thermalis* under far-red acclimation.

## 2. Materials and Methods

### 2.1 Cultivation conditions

*Choococcidiopsis thermalis* PCC7203, obtained from the Pasteur Culture Collection (PCC, France), was maintained at 30 °C under 10 µmol of photons m^-2^ s^-1^ of continuous white fluorescent light (L36W-840, OSRAM), in BG11 liquid medium (Rippka et al., 1979). To test the acclimation of PCC7203 to FR light, cultures in the exponential phase were resuspended in fresh BG11 medium and placed at 30 °C in orbital shaking at 100 rpm under 25 µmol of photons m^-2^ s^-1^ of respectively a simulated solar (SOL) and far red (FR) light. SOL light (Figure S1a) was simulated with the Star Light Simulator (SLS) extensively described in Claudi et al. (2021) and provided 20 µmol of photons m^-2^ s^-1^ of VIS light and 5 µmol of photons m^-2^ s^-1^ beyond 700 nm. The FR light was provided with a custom-made VIS-IR light simulator providing a 750 nm peaked light which resulted in 24,6 µmol of photons m^-2^ s^-1^ of FR light and only 0.4 µmol of photons m^-2^ s^-1^ of VIS light (Figure S1a). The distribution of light in the wavebands between 380 and 780 nm is reported in Figure S1b.

Cells growth during the acclimation was monitored via measurement of the optical density at 750 nm, OD_750_, as at this wavelength the absorption from pigments is negligible.

Once the acclimation to FR light was fully achieved, cultures were maintained in the exponential phase in the same conditions by weekly renewing them with fresh medium.

### 2.2 In vivo absorption spectra

To record *in vivo* absorption spectra, 2 ml of culture for each condition, were centrifuged at 1400 g for 10 min at room temperature. The pellet was homogenized with a pestle and 600 μl of fresh BG11 medium were added to solubilize the pellet. Samples were analyzed with a Cary100 UV-VIS spectrophotometer (Agilent) inside quartz cuvettes, exposing the opaque side to the light ray to minimize the scattering (Gan et al., 2014; Liistro et al., 2025).

### 2.3 Fluorescence analysis

Low temperature (77 K) fluorescence emission spectra were recorded using a Cary Eclipse Fluorescence Spectrometer (Agilent). Two milliliters of culture were centrifuged at 1500 g for 10 min at 4 °C and the pellet resuspended in 1 ml of glycerol 60% (w/v) and 10 mM Hepes pH 7.5. Samples were frozen in liquid nitrogen and stored in dark at −80 °C until analysis. For the analysis, samples were excited at 440 nm and emission spectra were recorded between 600 nm and 800 nm. Chlorophyll fluorescence and photosynthetic parameters *in vivo* were assessed with an AquaPen-C fluorometer (Photon Systems Instruments, Czech Republic). Prior to the analysis, cells were dark adapted for 3 minutes. Minimal fluorescence in dark adapted state, F_0_, was measured followed by a short saturating flash (620 nm, 1200 µmol of photons m^-2^s^-1^) to determine the maximal fluorescence in dark adapted state, F_m_. After a short dark relaxation, actinic light of 200 µmol of photons m^-2^s^-1^ was applied for 200 s and saturating flashes were delivered to measure the maximal fluorescence in light F_m_’. Minimal fluorescence in light, F_0_^’^, was also measured. Actinic light was then switched off and saturating pulses applied to measure maximal fluorescence in the dark after light exposure. The parameters of maximum quantum yield of PSII in dark-adapted state, F_v_/F_m_, effective quantum yield of PSII, Y(II), and non-photochemical quenching, NPQ, were calculated according to Genty et al., 1989; Bilger & Björkman, 1990; Kramer et al., 2004, as follows:

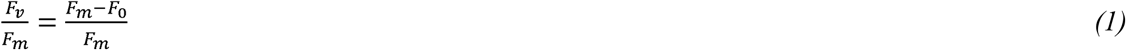

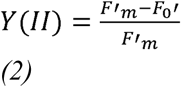

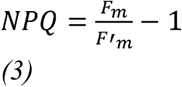

Analysis was performed on 4 ml of sample, with a chlorophyll concentration of approximately 1.5 µg per ml.

### 2.4 Pigment extraction and analysis

Lipophilic pigments were extracted by centrifuging cells for 10 min at 10000 g and resuspending the pellet in 1 ml of N,*N*-Dimethylformamide (DMF). Samples were kept in the dark at 4 °C for at least one day before analysis. For the extraction of hydrophilic pigments, cells were centrifuged for 10 min at 17500 g at 4 °C and to the pellet were added an equal volume of 150-212 µm acid washed glass beads (SIGMA) and 50 µl of phosphate buffer (Na_2_HPO_4_ 0.01 M, NaCl 0.15 M). Three cycles of bead beating were executed with a Bullet Blender Storm Pro Homogenizer (Next Advance) for 15 s at maximum speed alternated with 30 s in ice. 150 µl of phosphate buffer were added to the lysate and a last cycle of rupture was executed. Samples were centrifuged at 20000 g for 10 min at 4 °C and transferred to a new tube. 200 µl of phosphate buffer were added to the pellet and, after vortexing for resuspension, another centrifugation was performed. The supernatant was pooled with the pigment extract previously obtained and kept in ice; this last step was repeated until the supernatant appeared transparent, indicating the full extraction of hydrophilic pigments. Samples were kept in dark at −20 °C until analysis.

For the quantification of Chl *f*, pigments were extracted in 100% methanol, following the same protocol of cell rupture and extraction described for hydrophilic pigments.

Pigment extracts were analysed with a Cary100 UV-VIS spectrophotometer (Agilent). Concentration of chlorophyll *a* and total carotenoids were obtained with the Moran equation for DMF (Moran, 1982). Concentration of phycocyanin (PC) and allophycocyanin (APC) were obtained with Bennet and Bogorad equations (Bennett & Bogorad, 1973). Chl *f* was quantified with the following equation (Li et al., 2012):

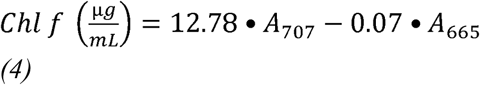

### 2.5 High Performance Liquid Chromatography (HPLC) coupled with UV/Vis spectroscopy and mass spectrometry (MS)

For HPLC analysis, cells were centrifuged at 17500 g for 10 min at 4 °C and equal volume of 150-212 µm acid washed glass beads (SIGMA) and 50 µl of acetone 90% were added to the pellet. Three cycles of bead beating were executed with a Bullet Blender Storm Pro Homogenizer (Next Advance) for 15 s at maximum speed alternated with 30 s in ice. 150 µl of acetone 90% were added to the lysate and a last cycle of rupture was executed. Samples were centrifuged at 20000 g for 10 min at 4 °C and transferred to a new tube. 200 µl of acetone 90% were added to the pellet and, after vortexing for resuspension, another centrifugation was performed. The supernatant was pooled with the extract previously obtained and kept in ice; this last step was repeated until the supernatant appeared transparent, indicating the full extraction of pigments. Analyses were performed with an Agilent 1100 series LC with a Lichrospher 100 RP column (250 mm length, 4 mm diameter) containing 5 μm silica particles coated by C-18 atom chains (Merck) as the stationary phase. The mobile phase consisted in two solutions: buffer A (methanol:acetonitrile:water in ratio 42:33:25) and buffer B (methanol: acetonitrile:ethyl acetate in ratio 50:20:30). The two solutions were eluted in the column according to the protocol optimized by Gan and colleagues (Gan et al., 2014) for Chl *d* and *f* detection, setting the Diode Array Detector (DAD) at 705 nm.

Selected samples were also analyzed with a mass spectrometer detector (MSD SL Trap), connected with the HPLC system. The ionization was performed within an atmospheric pressure chemical ionization source (APCI) in positive polarity with the following parameters: APCI temperature 350°C, nebulizer 60 psi, dry gas flow rate 12 L min^-1^, dry gas temperature 350°C, capillary voltage 3.5 kV, capillary exit 166.0 V, and skimmer 40 V.

### 2.6 Optically Detected Magnetic Resonance spectroscopy (ODMR) for triplet state detection

The principles of triplet-state ODMR are well established (Carbonera, 2009; Clarke, 1982). Briefly, pigment triplet states are generated under continuous illumination at low temperature while high-power microwaves drive magnetic transitions between triplet sublevels. Sweeping the microwave frequency across the region of expected Chl and Car triplet transitions, while simultaneously monitoring the optical response, induces resonant changes in absorption or fluorescence that link the optical and magnetic properties of the pigments. Fluorescence-Detected Magnetic Resonance (FDMR) reports chlorophyll triplet states through microwave-induced modulations of Chl fluorescence. Because carotenoid triplets arise via Chl→Car triplet transfer and carotenoids are essentially non-fluorescent, Car triplet transitions are detected indirectly through their signatures in Chl fluorescence. A more extensive description of the principles of the ODMR technique, specifically for carotenoid triplet detection, can be found in Figure S2 and S3.

ODMR spectroscopy was performed on a home-built spectrometer previously described in detail (Carbonera, 2009; Santabarbara et al., 2007; Carbonera et al., 1992). Light from a 250 W tungsten lamp is focused through a series of lenses on a 5 cm CuSO_4_ 0.6 M filter and subsequently on a 1.2 mm flat cell, where the sample is inserted. The flat cell is tilted so that its normal axis creates a 45° angle with respect to the direction of the incident light. Sample fluorescence emission, selected by long-pass filters (λ_fluo_ > 680 nm or λ_fluo_ > 715 nm) or by interferential filters (FWHM 10 nm) is collected in a standard 90° geometry by a photodiode (Centronic OSI 5K). A sweep oscillator (HP 8350B) generates microwaves that are amplified by a TWT amplifier (Sco-Nucletudes 10-46-30) and on/off amplitude modulated by a square wave pulse generator (Nuova Elettronica). Microwaves are sent to a slow-pitch helix in which the flat cell containing the sample is mounted. The cell is inserted in a liquid helium cryostat (Oxford instruments) pumped to reach a temperature of 1.8 K. The signals demodulated and amplified by a lock-in amplifier (EG&G model 5210) are reported as ΔF/F, where ΔF is the fluorescence intensity variation induced by the resonant microwaves and F is the steady state fluorescence intensity. Sample preparation consisted in diluting the *C. thermalis* PCC 7203 cell suspension in degassed glycerol (66% v/v) reaching an optical density of ∼ 1.0 at 750 nm. The resulting suspension was then introduced in the flat cell and rapidly frozen in the precooled cryostat. The presented data are the average of two different biological replicates.

### 2.7 Reactive oxygen species detection

To detect *in vivo* the presence of reactive oxygen species (ROS) in SOL and FR acclimated cells, 10 ml of the cultures were transferred into T-25 flasks and exposed to high white light (500 µmol of photons m^-2^ s^-1^) for 1 h in order to induce light stress and ROS production. Samples were then pelleted at 4000 g for 10 min at room temperature, resuspended in 1 ml pH 7 phosphate buffer (K_2_HPO_4_ 0.125 M, KH_2_PO_4_ 0.125 M) to which was added 2.5 µl of 2 mM 2’,7’-dichlorodihydrofluorescein diacetate (DCFH-DA) and finally incubated at room temperature for 1 h shaking in the dark. DCFH-DA serves as a fluorescent probe for oxidant detection. It is retained in cells in its non-fluorescent form (DCFH), but upon oxidation it is converted to 2′,7′-dichlorofluorescein (DCF), which emits fluorescence (Rajneesh et al., 2017; Singh et al., 2014).

The presence of fluorescent DCF inside PCC7203 cells was visualized using a confocal microscope (Leica SP5, Leica). Excitation was provided by a 405 nm laser diode (chlorophyll excitation) and a 488 nm Argon laser (probe excitation). Fluorescence signals were collected in two separate channels: 520–530 nm (DCF) and 672–689 nm (Chl a). Micrographs were analysed using FIJI-ImageJ software (National Institutes of Health, Bethesda, MD, USA), measuring the DCF fluorescence percentage on chlorophyll autofluorescence, providing an estimate of oxidative stress levels in living cells.

### 2.8 Oxygen production

To assess the photosynthetic efficiency and oxygen production rate of SOL and FR acclimated cells, 70 ml of culture with initial OD_750_ 0.6 were placed for two days in Atmospheric Simulation Chambers (ASCs), deeply described in (Battistuzzi et al., 2023). Briefly, ASCs are closed growth chambers that allow the cultivation of microorganisms under a desired light spectrum while constantly monitoring the CO_2_ and O_2_ concentration in the internal environment using two CO_2_ sensors (CO2M-20 and CO2M-100, SST sensing) and one O_2_ sensor (LOX-O2-S, SST sensing). The ASCs were maintained at 30 °C and the initial atmosphere supplied to the cells was ambient air enriched with 5% CO_2_. During these experiments the light intensity was maintained at 25 µmol photons m^-2^ s^-1^ for all tested spectra. Raw data of CO_2_ and O_2_ concentrations in ppm were converted in µmol by applying the law of ideal gasses and analysed through MatLab R2018b (MathWorks).

### 2.9 Statistical analysis

Statistical analyses were performed using the software GraphPad Prism v10.1.0, calculating mean and standard deviation for at least 3 biological replicates for each condition and comparing SOL and FR with ANOVA analysis (one-way or two-way depending on the dataset) and Tukey’s multiple comparison tests.

## 3. Results

### 3.1 Carotenoid content and composition changes upon FaRLiP

In order to assess the changes in carotenoid composition upon full achievement of far-red light photoacclimation (FaRLiP) with respect to cells acclimated to simulated solar light, *C. thermalis* cultures at initial OD_750_ of 0.25 were exposed to 25 µmol of photons m^-2^s^-1^ of respectively solar (SOL) and far-red (FR) light. As expected, *C. thermalis* cells under FR light grew more slowly than those under SOL light (Figure 1a). After 20 days, the mean OD_750_ reached 2.25 in SOL cultures but only 0.90 in FR-acclimating cultures. The development of the FaRLiP response in FR-acclimating cells was tracked by recording *in vivo* absorption spectra at days 3, 6, 9, 12, and 21 (Figure 1b). A characteristic absorption shoulder beyond 700 nm, indicative of FaRLiP induction, appeared as early as day 3 and progressively increased, reaching a maximum at day 21. After 21 days of FR light exposure, the absorption level beyond 700 nm matches the one from fully acclimated cultures, constantly maintained in monochromatic FR light for two months, indicating that cells were completely acclimated. The red-shifted absorption is attributable to the synthesis of FR-absorbing pigments, namely Chl *d*, Chl *f*, and FR-APC. To confirm whether the red-shift originated from both chlorophylls and FR-APC, absorption spectra of pigment extracts were also analyzed. Indeed, both lipophilic and hydrophilic pigment extracts presented shoulders of absorption beyond 700 nm (Figure S4a, S4b). Consistently, low temperature (77K) fluorescence measurements registered at day 21 highlighted the acclimation to FR light in C. *thermalis* cells (Figure S4c). The maximum fluorescence emission in SOL-acclimated cells was at 724 nm, corresponding to white-light PSI, while in FR-acclimated cells the maximum emission shifted to 752 nm, indicating the presence of FR-PSs.

**Figure 1.**
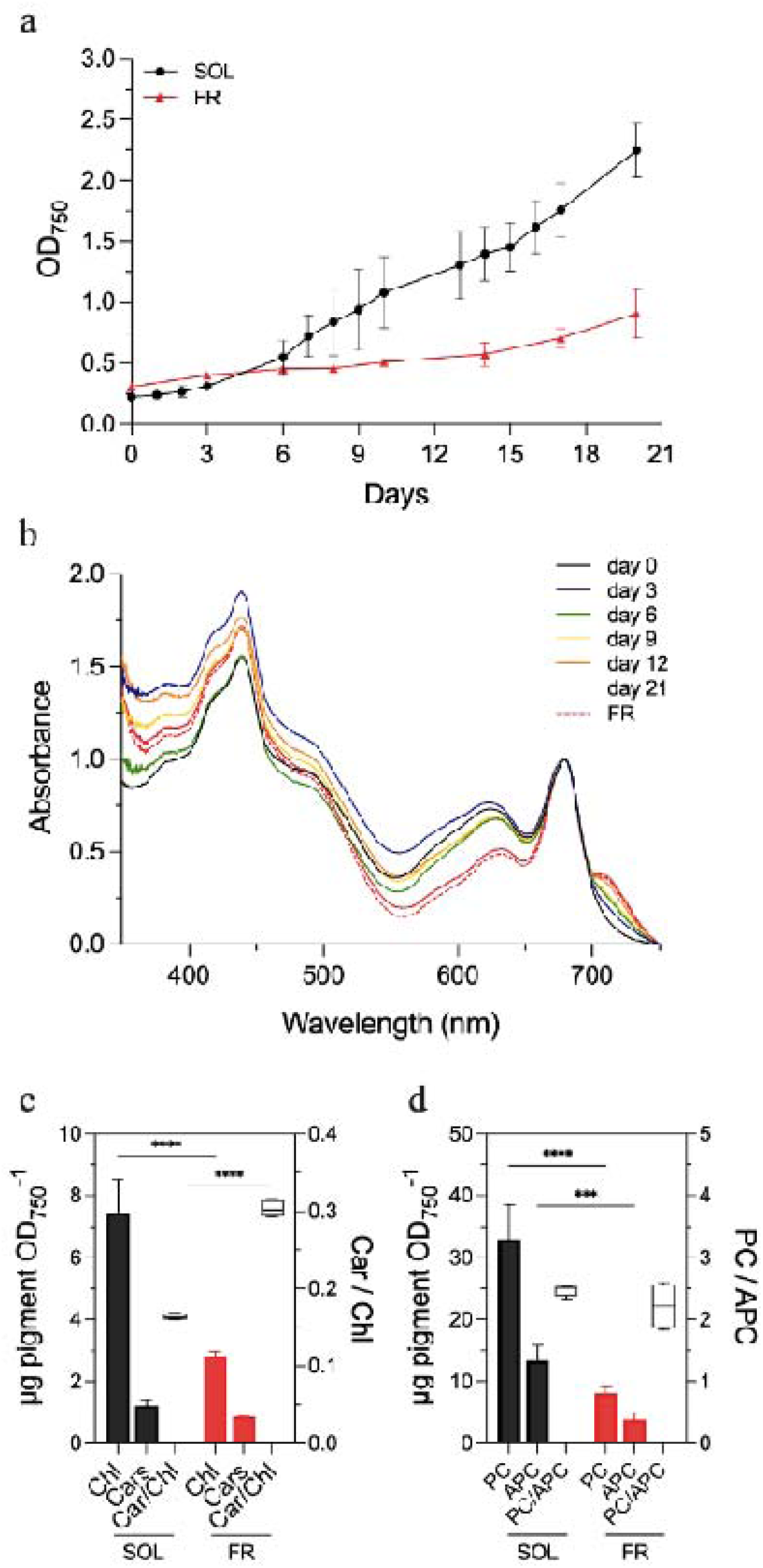
Acclimation of *C. thermalis* PCC7203 to SOL and FR light. a) culture OD_750_ during the 21 days of acclimation to SOL and FR light; b) *in vivo* absorption spectra of FR-acclimating cells during the 21 days of FR light exposure and fully acclimated (red dashed line) normalized at 680 nm; c) chlorophyll and carotenoid content of cells at day 21 of SOL and FR light exposure expressed as µg of pigment per OD_750_ (left axis) and carotenoid to chlorophyll ratio (right axis), two-way ANOVA, p-value <0.0001; d) phycobiliprotein content of cells at day 21 of SOL and FR light exposure expressed as µg of pigment per OD_750_ (left axis) and phycocyanin to allophycocyanin ratio (right axis) two-way ANOVA, ****: p-value <0.0001, ***: p-value =0.0002. SOL: solar light-acclimating cells, FR: far red light-acclimating cells, Chl: chlorophyll *a*, Cars: carotenoids, PC: phycocyanin, APC: allophycocyanin.

Pigment content analysis further revealed a reduction in total pigments – both chlorophylls and phycobiliproteins (PBPs) – in FR-acclimated cells compared with SOL-acclimated cells (Figure 1c, 1d). In contrast, carotenoid levels were not significantly reduced. Indeed, carotenoids to chlorophyll ratio increased from 0.16 in SOL-acclimated cells to 0.30 in FR-acclimated ones.

Quantification of Chl *f* in methanol extracts showed that FR-acclimated cells contained 2.83 µg Chl *a* per OD_750_ and an additional 0.31 µg Chl *f* per OD_750_, corresponding to ∼10% of the total chlorophyll content. Moreover, to confirm the presence of both Chl *d* and *f* and to verify if carotenoids composition changed after full FaRLiP acclimation, reverse-phase liquid chromatography was performed (Figure 2a). Chromatograms and DAD spectra highlighted the presence of Chl *d* and *f* in FR-acclimated cells, which were absent in SOL-acclimated cells (Figure S5). Carotenoids were identified based on DAD spectra and retention times, supported by mass spectrometry for ambiguous peaks (canthaxanthin and echinenone), which confirmed the predicted assignments (Figure 2, S6, S7). Detected carotenoids included myxol-2′-fucoside, canthaxanthin, echinenone, β-carotene, and cis-β-carotene, all present in both SOL- and FR-acclimated cells.

**Figure 2.**
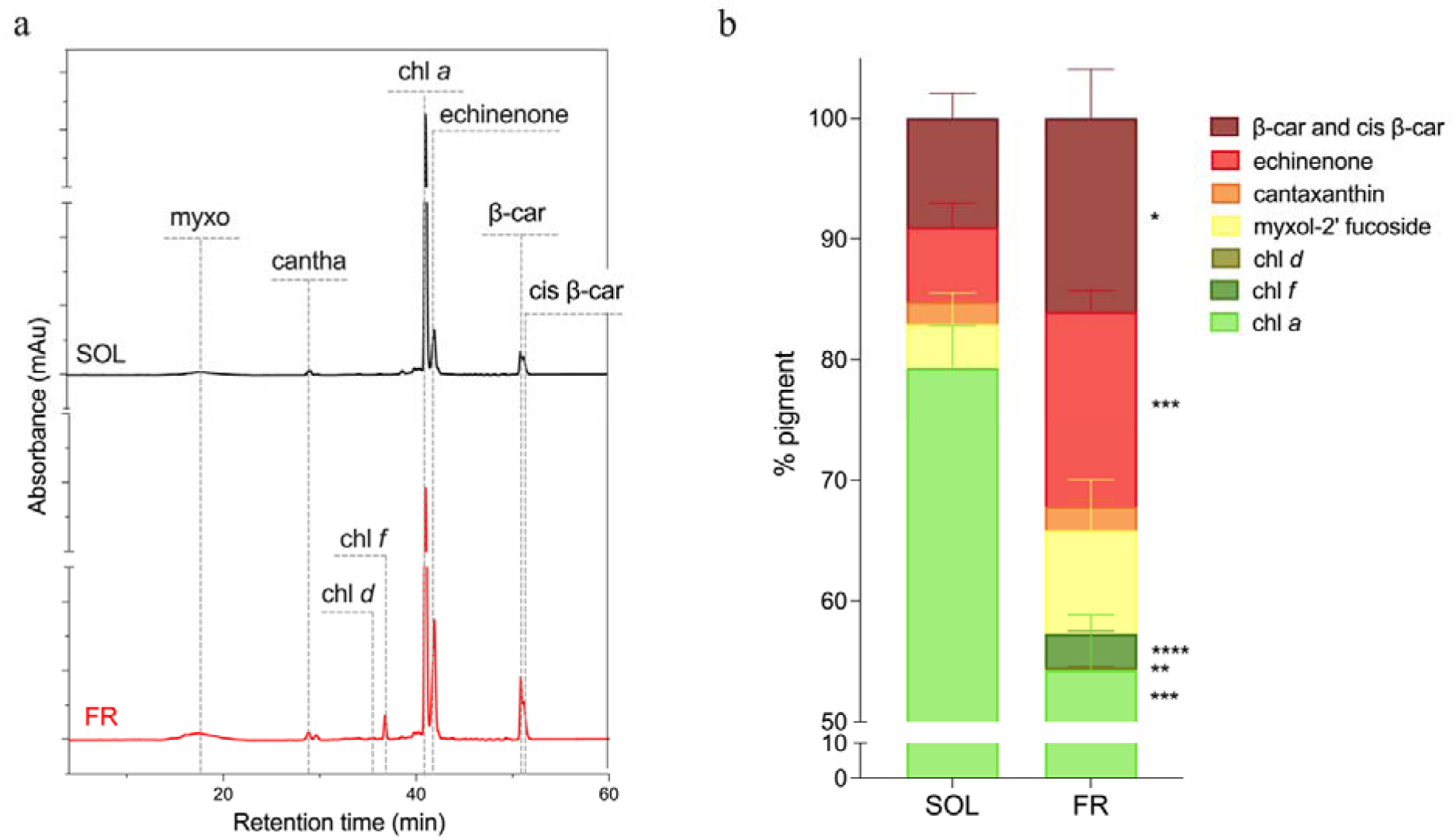
Chlorophyll and carotenoids composition of *C. thermalis* cells acclimated to SOL and FR light. a) HPLC chromatograms at 440 nm of extracts from cells acclimated to SOL (black) and FR (red) light, dotted lines indicate the retention times of the detected pigments; b) relative quantification of the detected pigments, expressed as percentage of the total integrated area of absorbance in the chromatograms. For b) unpaired t test was executed for each pigment in SOL and FR: β-carotene and cis β-carotene (*) p-val = 0.0431, echinenone (***) p-val = 0.0009, chl *f* (****) p-val < 0.0001, chl *d* (**) p-val = 0.0019, chl *a* (***) p-val = 0.0005. Myxo: Myxol-2’ fucoside; cantha: canthaxanthin; chl *d*: chlorophyll *d*; chl *f*: chlorophyll *f*; chl *a*: chlorophyll *a*; β-car: β-carotene; cis β-car: cis β-carotene; SOL: solar; FR: far-red.

Relative quantification of pigments based on the comparison of integrated areas of absorbance in chromatograms revealed the strong decrease of chlorophylls percentage in FR light, together with the concurrent rise in carotenoids abundance (Figure 2b). However, changes among individual carotenoids were not uniform upon acclimation to FR light. Myxol-2’ fucoside increased from 3.6% in SOL to 8.6% in FR, canthaxanthin slightly increased from 1.8% to 2.0%. Echinenone and β-carotene (comprehensive of cis-β-carotene) significatively increased from 6.2% and 9.0% in SOL to 16.1% and 16.0% in FR, respectively.

### 3.2 ^3^Chl and ^3^Car levels

To detect and characterize possible harmful photogeneration of ROS-producing chlorophyll triplet states and consequently assess the differences in photoprotection activities occurring in C. *thermalis* cells acclimated to SOL and FR luminous exposures, *in cellula* Optically Detected Magnetic Resonance (ODMR) was performed. The technique makes it possible to reveal the formation of chlorophyll triplet states (^3^Chl) bound to either PSII or PSI, that represent a dangerous source of singlet oxygen leading to oxidative stress. In photosynthetic systems, chlorophyll triplets can be quenched by transfer to an adjacent carotenoid, if the geometrical requirements of sufficient proximity and correct orientation between the two molecules are fulfilled. Carotenoid triplet states are not able to sensitize singlet oxygen formation, and they dissipate the excess energy, performing in this way a photoprotective function (Di Valentin & Carbonera, 2017; J. Ho et al., 2017).

ODMR spectra for *C. thermalis* SOL and FR cells are reported in Figure 3. Spectra have been collected by looking at the total fluorescence emission from the two different samples. Total fluorescence was collected at λ_fluo_ > 680 nm for SOL cells and at λ_fluo_ > 715 nm for FR cells, given that the latter exhibits a markedly red-shifted fluorescence at low temperatures.

**Figure 3.**
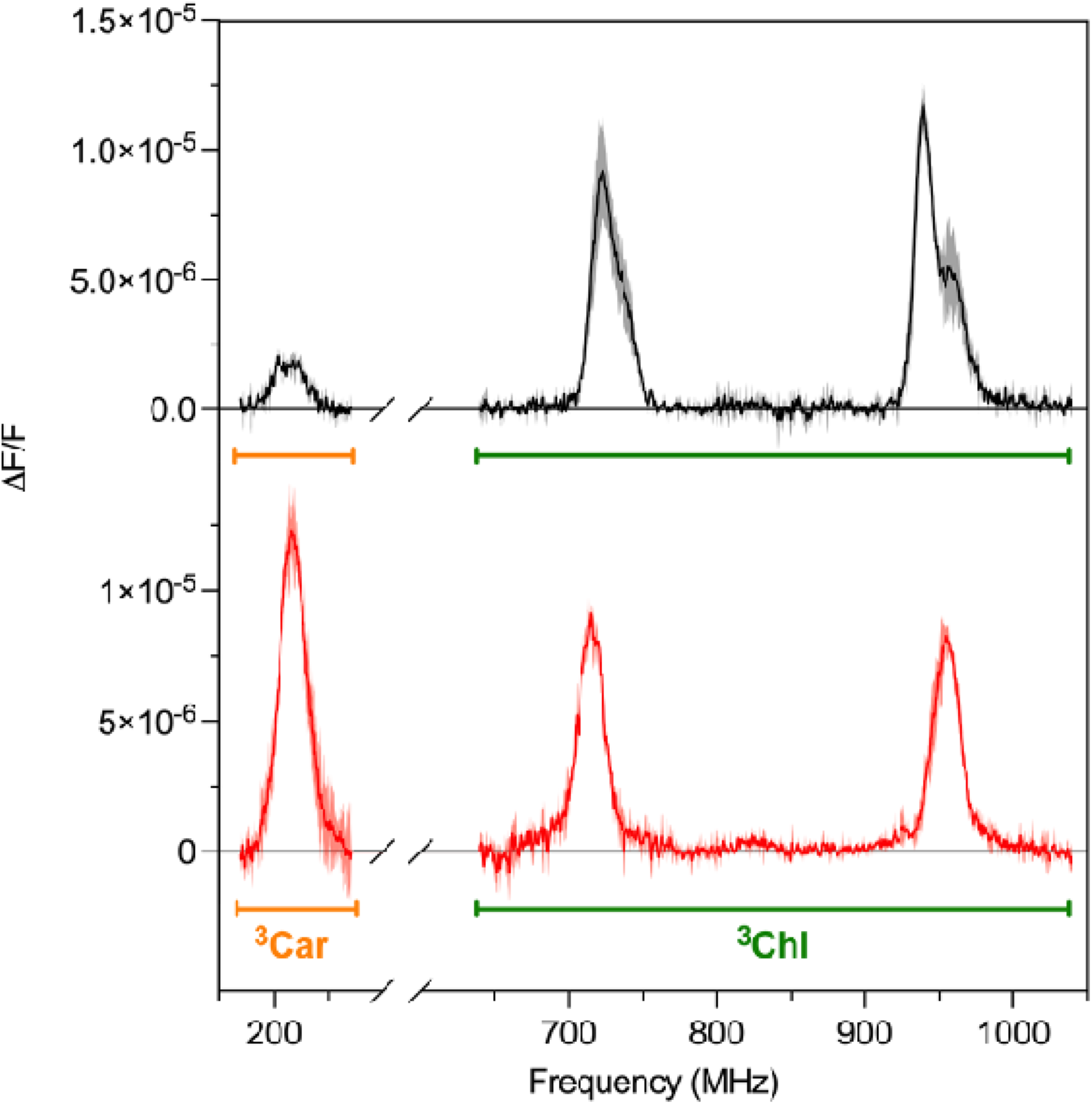
FDMR spectra of *C. thermalis* cells acclimated to SOL and FR light. Fluorescence detected magnetic resonance of ^3^Chl (on the |D|-|E| and |D| + |E| transitions underlined by the green bar) and ^3^Car (2|E| transition, orange bar) for both SOL (top black trace) and FR (bottom red trace) acclimated cells. FDMR signals detected by collection of the total sample fluorescence emission through long-pass filters, specifically _fluo_> 680 nm for SOL cells and _fluo_> 715 nm for FR cells. SOL and FR spectra have been vertically shifted for clarity. Microwave on/off modulation frequency of 33 Hz (for ^3^Chl) or 333 Hz (for ^3^Car), temperature of 1.8 K.

For SOL acclimated cells, ^3^Chl transitions bands are observed in the frequency ranges of 680–760 MHz and of 900–980 MHz (Figure 3, black trace), respectively belonging to the |D| - |E| and |D| + |E| triplet transitions of Chl *a*. Both bands are split into two components: the first one at 722/940 MHz and the second at 739/958 MHz. Comparisons of these values with already published data, for example from *Chlamydomonas reinhardtii* (Santabarbara et al., 2007), indicate that the signals may originate mainly from unquenched ^3^Chl *a* belonging to PSI. At lower frequencies, where the 2|E| transition of *^3^*Car is expected, a positive band centered at 211 MHz is exhibited, showing that carotenoids are active in photoprotection.

FDMR spectrum of FR cells (Figure 3, red trace) in the ^3^Chl region shows two positive bands centered at 715 MHz and at 956 MHz. Attribution of these features to ^3^Chl *a* is rather straightforward considering that they are positioned at a very similar frequency with respect to the SOL sample. |D| - |E| and |D| + |E| transitions of ^3^Chl *a* in FR cells appear to be constituted of just one single component. No ^3^Chl *d* or ^3^Chl *f* were detected in the FR cells FDMR, which is a surprising outcome since low energy pigments are expected to act as final energy acceptors and constitute a site of formation of triplet states, especially at the very low temperature of the experiments.

Looking instead at the ^3^Car triplets, an intense positive band centered at 216 MHz reveals the presence of a remarkable photoprotective activity in the FR acclimated cells, evidenced by the stronger intensity of the ^3^Car transition compared to the corresponding one for SOL cells. The ^3^Car/^3^Chl ratio (area of the 2|E| ^3^Car band compared to the |D|+|E| ^3^Chl *a* band) was 1.82 for FR cells and 0.25 for SOL cells.

Figure 4a reports FDMR spectra of FR cells selectively detected at wavelengths from 730 nm to 770 nm, together with a plot of their integrated intensities (Figure 4b). Transitions started to show at _fluo_ = 740 nm and increased in intensity until a maximum was reached between 750 and 760 nm. The wavelength of the maximum FDMR signal matches the peak of low temperature fluorescence emission of FR-acclimated cells (Figure S4c), suggesting photoprotection coupling with the main photosystem’s emitters, i.e. far-red shifted chlorophylls

**Figure 4.**
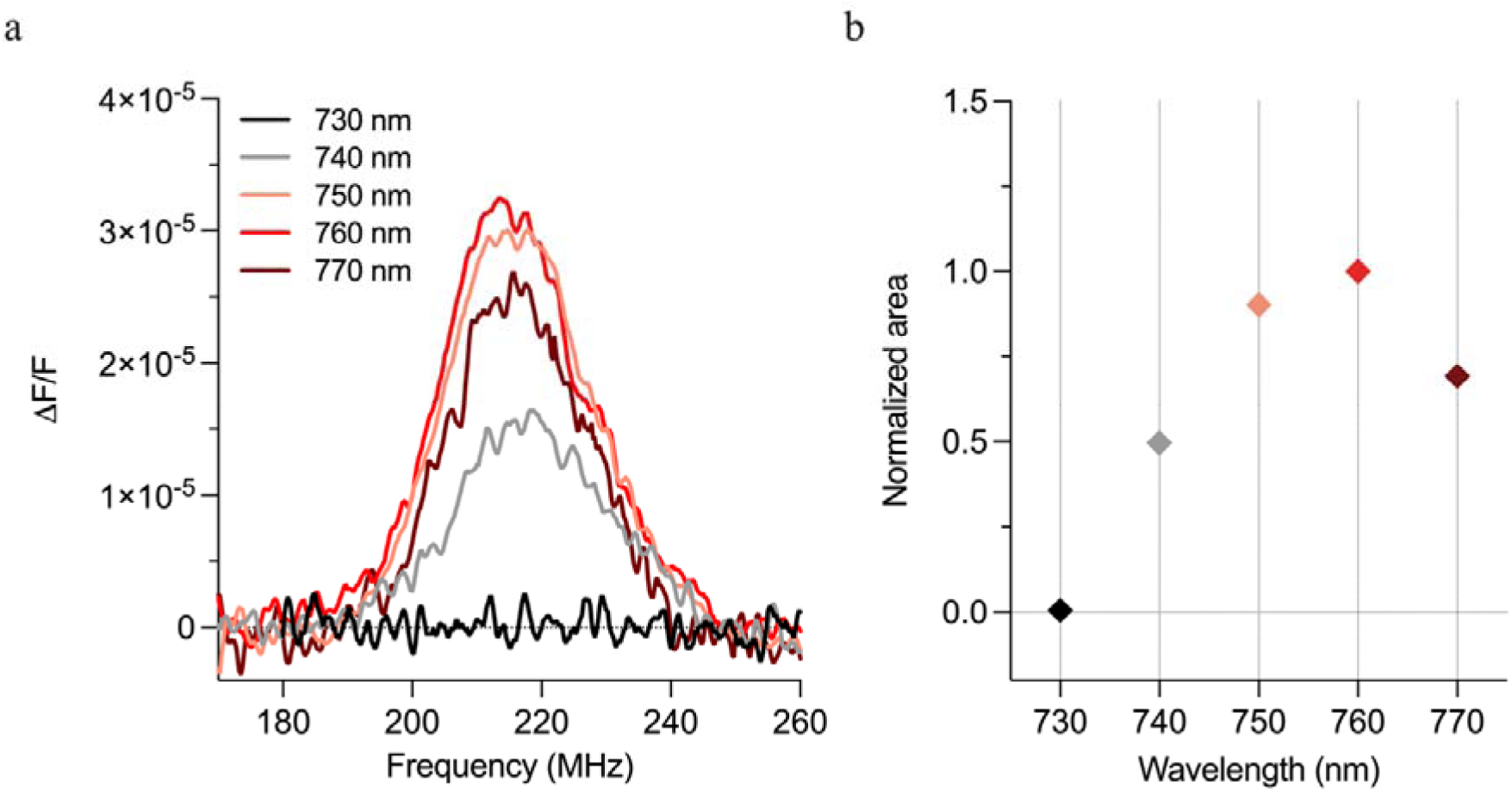
^3^Car FDMR spectra of *C. thermalis* FR cells at specific emission wavelengths. a) Fluorescence detected magnetic resonance spectra of ^3^Car (on the 2|E| transition) recorded at specific wavelengths through interferential filters for FR acclimated cells. b) Area of the transition bands as a function of the fluorescence detection wavelengths. The area of the most intense band was normalized to one. Microwave on/off modulation of 333 Hz (for ^3^Car), temperature of 1.8 K. Error is estimated to be around 10 % for each value of integrated area.

### 3.3 NPQ and ROS production

Photosynthetic parameters like PSII quantum yield and NPQ were measured *in vivo* on SOL- and FR-acclimated cells (Figure 5).

**Figure 5.**
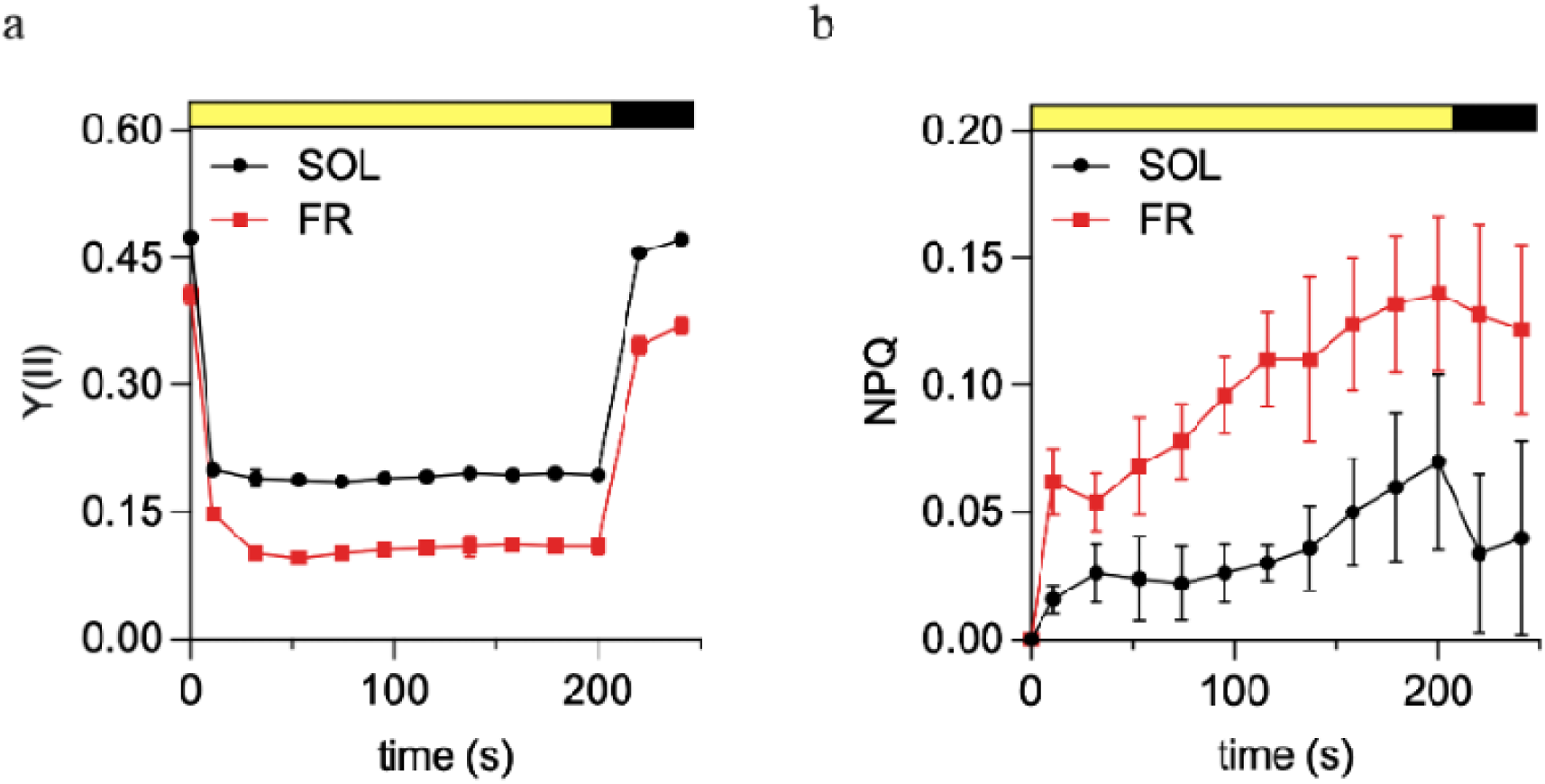
*In vivo* chlorophyll fluorescence analysis of *C. thermalis* cells acclimated to SOL and FR light. Dark adapted cells were exposed for 200 s to 200 µmol of photons m^-2^s^-1^ light, followed by 60 s of darkness. Quantum yield of Photosystem II, Y(II), (a) and non-photochemical quenching, NPQ, (b) of SOL-(black) and FR-(red) acclimated cells are shown.

The maximum photochemical quantum yield of PSII was significantly higher (p-val <0.0001) in SOL (0.47) than in FR (0.41). This difference, though, is likely influenced by the diverse content in phycobiliprotein, that is lower in cells exposed to FR light. However, the kinetics of the effective quantum yield were similar in the two conditions, indicating a comparable functionality of PSII upon illumination. On the other hand, FR-acclimated cells showed a higher non-photochemical quenching than SOL-acclimated, also with a faster activation, suggesting a stronger photoprotective response following the acclimation to longer wavelengths, even though the values reached are rather low in both conditions.

The presence of reactive oxygen species, ROS, was detected using the oxidant-sensing probe 2’,7’-dichlorodihydrofluorescein diacetate, DCFH-DA, which is retained in cells in the non-fluorescent DCFH. DCFH is oxidized by ROS in 2’,7’-dichlorofluorescein, emitting a cyan-green fluorescence (520-530 nm), well distinguishable from chlorophyll fluorescence. After a high-light treatment of 500 µmol of photons m^-2^ s^-1^ for 1 hour, cells incubated with DCFH-DA were imaged for DCF fluorescence and chlorophyll *a* fluorescence. Results show a higher percentage of DCF fluorescence in SOL acclimated samples (Figure 6), indicating that FR acclimated cells experienced a less oxidative stress, facing the high-light treatment more efficiently.

**Figure 6.**
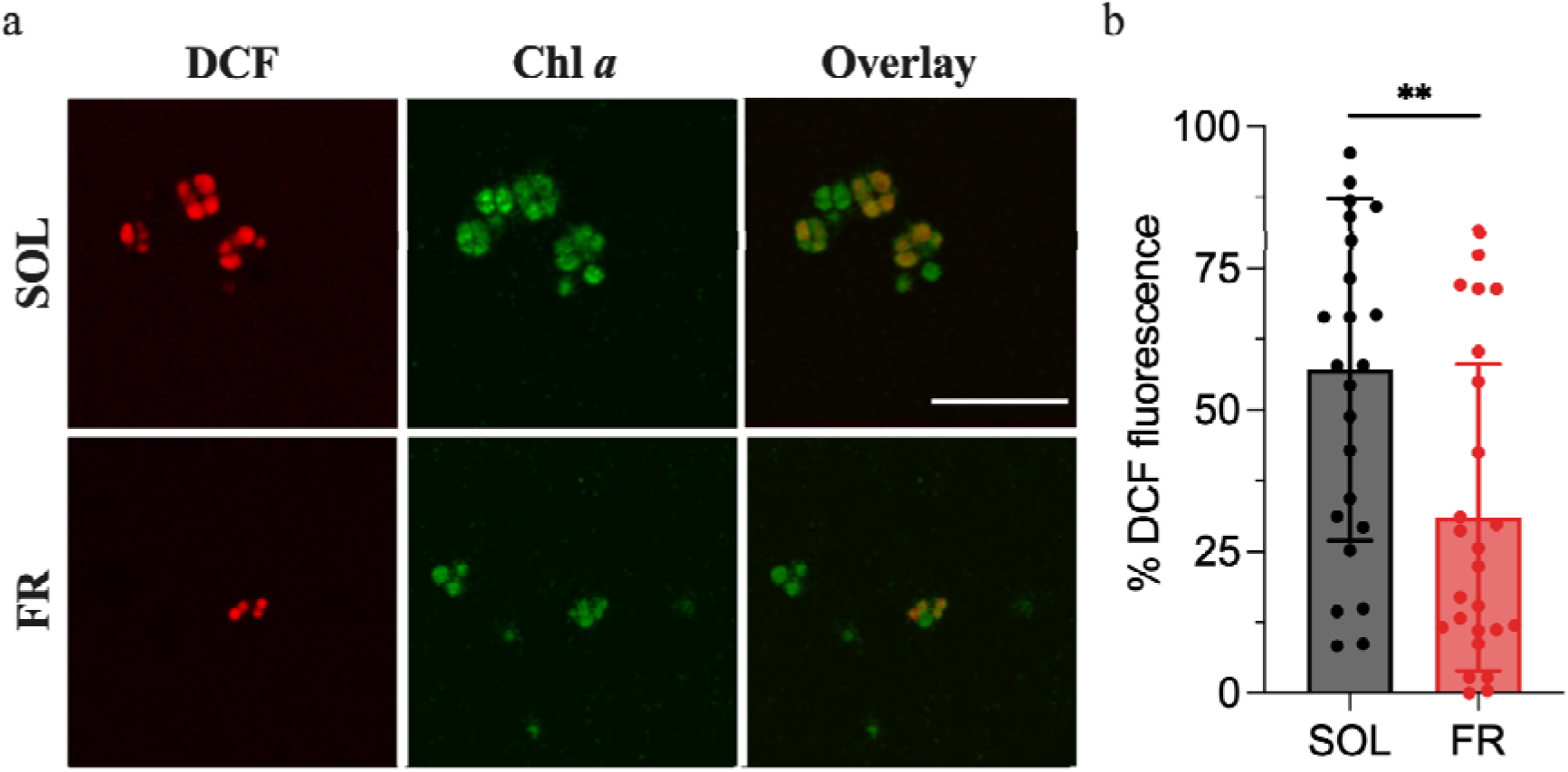
Reactive oxygen species in *C. thermalis* cells acclimated to SOL and FR light treated with DCF. a) Fluorescence emission registered with confocal microscopy in SOL and FR acclimated cells stressed with high light of DCF (proxy for oxidative stress, red), Chl *a* (proxy for cell vitality, green), and overlay of both channels, scale bar correspond to 25 µm; b) percentage of DCF fluorescence on chlorophyll *a* fluorescence unpaired t test on <26 micrographs, p-value <0.005. DCF: 2’,7’-dichlorofluorescein; Chl *a*: chlorophyll *a*; SOL: solar; FR: far red.

Given the greater protection against ROS formation observed in FR-acclimated cells after exposure to stressing white light, as seen by the screening performed with the fluorescent probe, we investigated whether a sudden interaction with SOL light of FR-acclimated cells would lead to increased O production, potentially explaining their capability to enhance resistance to oxidative stress. Cells were placed in the atmospheric simulating chambers, ASCs, and supplied with ambient air enriched with 5% CO_2_ in order to evaluate the O_2_ released in the closed atmosphere and the O_2_ production rate (Figure 7). We tested both FR and SOL light acclimated cultures, to evaluate their O_2_ production capacity under the respective light of growth. As expected, considering the difference in growth capability (Figure 1), FR acclimated samples exposed to FR light produced a small amount of O_2_, with a constant low rate. On the other hand, when FR acclimated cells were suddenly exposed to SOL light, they demonstrated a noticeable release of O_2_, with an increase in the production rate higher than the one exhibited by SOL acclimated cells. The FaRLiP photosynthetic apparatus is still able to sensibly photosynthesize in visible light and generate O_2_, progressively increasing its release since the start of the exposure.

**Figure 7.**
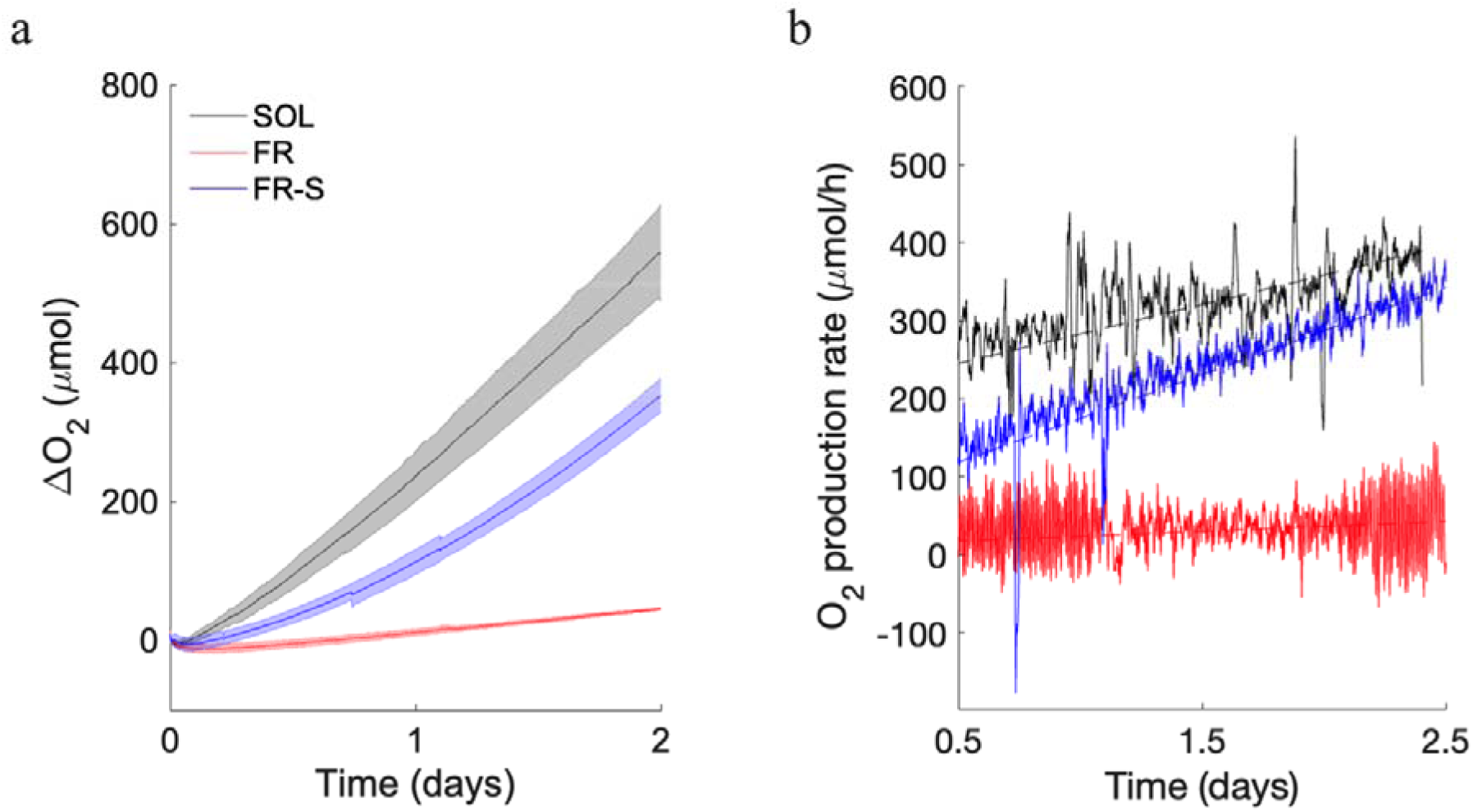
O_2_ evolution of *C. thermalis* cells acclimated to SOL and FR light. a) O_2_ evolution in an initially anoxic atmosphere by SOL acclimated cells placed under SOL light (black line) and FR acclimated cells placed either in FR light (red line) and SOL light (blue line), expressed as Δµmol of O_2_; b) rates of O_2_ evolution (solid lines), expressed as µmol of gas produced per day, dotted lines indicate rates’ linear regressions. SOL: solar; FR: far red; FR-S: far-red acclimated cells exposed to solar light.

## 4. Discussion

FaRLiP acclimation involves the expression of a gene cluster encoding red-shifted pigments and photosynthetic subunits that remodel the photosynthetic apparatus (Wolf & Blankenship, 2019; Gan et al., 2014). As a result, FaRLiP strains exposed to far-red light display a characteristic absorption shoulder beyond 700 nm. In several species (e.g., *Chroococcidiopsis thermalis* PCC7203, *Calothrix* sp. PCC7507, *Fischerella thermalis* PCC7521, *Chlorogleopsis* sp. PCC9212, and *Halomicronema hongdechloris*), this acclimation is also accompanied by enhanced absorption between 450–500 nm (Schmitt & Friedrich, 2024; Gan et al., 2015), which in *C. thermalis* has been attributed to an increased carotenoid content (Mascoli et al., 2022). However, the specific carotenoids involved, and their functional roles remain poorly understood.

In this work, we investigated the carotenoid composition of far-red acclimated *C. thermalis* cells, with a focus on photoprotection. In fully acclimated cells, the carotenoid-to-chlorophyll ratio doubled compared to SOL-acclimated cells. Since in the employed experimental setup the FR spectrum to which cells were acclimated supplies only 0.4 µmol photons m² s¹ of visible light, out of 25 µmol photons m² s¹, carotenoid-to-chlorophyll ratio increase is unlikely to functionally enhance light harvesting. Instead, we hypothesize it fulfills a photoprotective function.

Two key photoprotective carotenoids were found to increase: β-carotene and echinenone. β-carotene is known to quench both excited chlorophyll triplet state and singlet oxygen, thereby suppressing ROS formation (Fiebig et al., 2023; Hashimoto et al., 2016). Echinenone, for its part, participates in NPQ as a binding carotenoid of the orange carotenoid protein (OCP) and as the precursor of 3′-hydroxyechinenone, the most common OCP ligand (Niziński et al., 2025; Wilson et al., 2006; Kerfeld, 2004).

Direct quenching of ^3^Chls by β-carotene requires close spatial coupling with chlorophylls. Indeed, FDMR analysis revealed that FR-acclimated cells exhibited a markedly stronger ³Car signal than SOL-acclimated cells, with the ³Car/³Chl ratio increasing sevenfold. Strikingly, the wavelength dependence of the intensity of FDMR ³Car transitions (Figure 4b) overlapped with the 77 K fluorescence emission profile of FR cells (Figure S2c), suggesting that the carotenoids engaged in triplet quenching are close to the chlorophyll emitters. Since low-energy Chl *f* pigments dominate fluorescence in FR cells, they are likely to be more prone to intersystem crossing and thus at higher risk of triplet-induced photodamage. The observation that the ³Car FDMR signal peaks at the Chl *f* emission maximum, together with the absence of unquenched ³Chl *f* signals, suggests that FR-acclimated *C. thermalis* has evolved a pigment arrangement in which carotenoids specifically safeguard red-shifted chlorophylls from photodamage. In support of this hypothesis, an early cryo-EM structure of FR-PSI (Kato et al., 2020) has shown that there are indeed some Chl *f* – β-carotene couples that are particularly close in space (for example the one indicated as f825-BCR847B, Kato *et al*. notation).

Screening with the oxidant-sensing probe DCFH-DA confirmed that, under the same high-light stress, FR-acclimated cells generated significantly less ROS, indicative of a less oxidative intracellular environment. The Chl triplet quenching mediated by carotenoids reduces the probability of excited chlorophylls reacting with molecular oxygen, thereby preventing the formation of harmful ROS (Fiebig et al., 2023). The protection likely arises not only from chlorophyll–carotenoid coupling but also from the direct quenching of singlet oxygen (¹O) (Latifi et al., 2009), which may mainly occur at the level of thylakoidal membranes. Indeed, the higher carotenoid content encountered in FR cells, also associated with β-carotene, is reasonably involving a larger concentration of it inside the cell membranes, as structural data describing FR-PSI and FR-PSII chromophore-protein arrangements do not show major differences in the total number of bound pigments with respect to canonical photosystems.

A crucial role in maintaining a non-oxidative cell environment and preserving the functionality of the photosynthetic apparatus is also played by OCP, that in cyanobacteria mediates the fastest component of non-photochemical quenching (Arshad et al., 2022; Wilson et al., 2006) by dissipating into heat excessive light excitation absorbed by phycobilisomes. Quenching of singlet oxygen by direct interaction with the internal OCP’s carotenoid was also proposed as an alternative mechanism for OCP antioxidant activity (Sedoud et al., 2014). The OCP of *C. thermalis* PCC7203 belongs to the family called OCPX, generally associated with ancestral cyanobacteria (Bao et al., 2017). Little information is currently available for this highly heterogeneous clade, recently suggested to be further divided into three subgroups (Slonimskiy et al., 2022). Data obtained so far evidenced a similar capacity of OPCX to quench phycobilisome fluorescence compared to the conventional and more represented family of OCP, called OCP1, while the back conversion from its active to inactive state seems to be faster and less temperature dependent for *in vitro* OCPX (Slonimskiy et al., 2022; Muzzopappa et al., 2019). This was indeed proposed to advantage extremophile bacteria like *C. thermalis* (Slonimskiy et al., 2022). The experiments measuring fluorescence evolution in time show that FR-acclimated cells displayed both a faster activation and a higher overall NPQ capacity compared to SOL-acclimated cells. Nonetheless, the NPQ levels are relatively low in both conditions, suggesting a higher yet limited photoprotection depending on this mechanism in FR-acclimated cells. To our knowledge, no indications have been reported so far for specific interactions between FR-PBS and OCP (of any family), which may be worth exploring in the future.

Overall, *C. thermalis* cells exhibit a robust set of photoprotective mechanisms coupled to the FaRLiP response, an apparently counterintuitive feature, given the lower energy of far-red photons. However, when FR-acclimated cells were transferred to visible light, they started to quickly produce a sensible amount of O_2_, which, in the context of a sub-optimal apparatus in visible light, could represent a condition of higher stress than in cells presenting white-light photosystems.

Within the FaRLiP photosynthetic apparatus, where energy trapping and charge separation are slowed by red-shifted pigments, thereby reducing photosynthetic efficiency (Mascoli et al., 2020, 2022), effective photoprotection may be essential to prevent damage arising from intersystem crossing and triplet generation. Because of their weaker photosynthetic machinery, cyanobacteria acclimated to far-red light could be particularly vulnerable to oxidative stress upon sudden exposure to visible light. For those strains that inhabit niches interested by periodical or episodical variations in light quality, managing to proficiently dissipate excess energy through NPQ and Chl triplet quenching could ensure a safer transition to visible light and faster conversion to visible-like photosystems. A robust photoprotective capacity against high-energy short-wavelength radiation may be critical in photosynthetic organisms utilizing a FaRLiP apparatus not optimized for visible light.

Notably, as far-red photosynthesis emerged early in cyanobacterial evolution, the coupling of strong photoprotection with FR-driven photosystems may have been essential when Earth’s atmosphere lacked protective layers. Moreover, this strategy could have been advantageous under fluctuating irradiance conditions during the initial stages of terrestrial colonization.

## CRediT authorship contribution statement

EL: Formal analysis, Investigation, Methodology, Data curation, Visualization, Writing – original draft, Writing – review and editing; AC: Formal analysis, Investigation, Methodology, Data curation, Writing – original draft, Writing – review and editing; AA: Formal analysis, Investigation, Methodology, Data curation, Writing – review and editing; SP: Formal analysis, Methodology, Data curation, Writing – review and editing; EM: Formal analysis, Methodology, Data curation, Writing – review and editing; NLR: Resources, Conceptualization, Investigation, Supervision, Writing – original draft, Writing – review and editing; DC: Resources, Conceptualization, Investigation, Supervision, Writing – original draft, Writing – review and editing.

## References

Airs, R. L., Temperton, B., Sambles, C., Farnham, G., Skill, S. C., & Llewellyn, C. A. (2014). Chlorophyll *f* and chlorophyll *d* are produced in the cyanobacterium *Chlorogloeopsis fritschii* when cultured under natural light and near-infrared radiation. FEBS Letters, 588(20), 3770–3777. 10.1016/j.febslet.2014.08.026

Antonaru, L. A., Cardona, T., Larkum, A. W. D., & Nürnberg, D. J. (2020). Global distribution of a chlorophyll f cyanobacterial marker. ISME Journal, 14(9). 10.1038/s41396-020-0670-y

Antonaru, L. A., Rad-Menéndez, C., Mbedi, S., Sparmann, S., Pope, M., Oliver, T., Wu, S., Green, D. H., Gugger, M., & Nürnberg, D. J. (2025). Evolution of far-red light photoacclimation in cyanobacteria. Current Biology, 35(11), 2539–2553.e4. 10.1016/j.cub.2025.04.038

Arshad, R., Saccon, F., Bag, P., Biswas, A., Calvaruso, C., Bhatti, A. F., Grebe, S., Mascoli, V., Mahbub, M., Muzzopappa, F., Polyzois, A., Schiphorst, C., Sorrentino, M., Streckaité, S., van Amerongen, H., Aro, E.-M., Bassi, R., Boekema, E. J., Croce, R., … Büchel, C. (2022). A kaleidoscope of photosynthetic antenna proteins and their emerging roles. Plant Physiology, 189(3), 1204–1219. 10.1093/plphys/kiac175

Bao, H., Melnicki, M. R., Pawlowski, E. G., Sutter, M., Agostoni, M., Lechno-Yossef, S., Cai, F., Montgomery, B. L., & Kerfeld, C. A. (2017). Additional families of orange carotenoid proteins in the photoprotective system of cyanobacteria. Nature Plants, 3(8), 17089. 10.1038/nplants.2017.89

Battistuzzi, M., Cocola, L., Claudi, R., Pozzer, A. C., Segalla, A., Simionato, D., Morosinotto, T., Poletto, L., & La Rocca, N. (2023). Oxygenic photosynthetic responses of cyanobacteria exposed under an M-dwarf starlight simulator: Implications for exoplanet’s habitability. Frontiers in Plant Science, 14. 10.3389/fpls.2023.1070359

Bennett, A., & Bogorad, L. (1973). Complementary chromatic adaptation in a filamentous blue-green alga. The Journal of Cell Biology, 58(2), 419–435. 10.1083/jcb.58.2.419

Bilger, W., & Björkman, O. (1990). Role of the xanthophyll cycle in photoprotection elucidated by measurements of light-induced absorbance changes, fluorescence and photosynthesis in leaves of Hedera canariensis. Photosynthesis Research, 25(3), 173–185. 10.1007/BF00033159

Billi, D., Napoli, A., Mosca, C., Fagliarone, C., de Carolis, R., Balbi, A., Scanu, M., Selinger, V. M., Antonaru, L. A., & Nürnberg, D. J. (2022). Identification of far-red light acclimation in an endolithic Chroococcidiopsis strain and associated genomic features: Implications for oxygenic photosynthesis on exoplanets. Frontiers in Microbiology, 13. 10.3389/fmicb.2022.933404

Carbonera, D. (2009). Optically detected magnetic resonance (ODMR) of photoexcited triplet states. Photosynthesis Research, 102(2–3), 403–414. 10.1007/s11120-009-9407-5

Carbonera, D., Giacometti, G., & Agostini, G. (1992). FDMR of Carotenoid and Chlorophyll triplets in light-harvesting complex LHCII of spinach. Applied Magnetic Resonance, 3(5), 859–872. 10.1007/BF03260117

Chen, M., Li, Y., Birch, D., & Willows, R. D. (2012). A cyanobacterium that contains chlorophyll *f* – a red[absorbing photopigment. FEBS Letters, 586(19), 3249–3254. 10.1016/j.febslet.2012.06.045

Clarke, R. H. (1982). Triplet State ODMR Spectroscopy: Techniques and Applications to Biophysical Systems. Wiley.

Claudi, R., Alei, E., Battistuzzi, M., Cocola, L., Erculiani, M. S., Pozzer, A. C., Salasnich, B., Simionato, D., Squicciarini, V., Poletto, L., & La Rocca, N. (2021). Super-earths, m dwarfs, and photosynthetic organisms: Habitability in the lab. Life, 11(1). 10.3390/life11010010

Consoli, G., Tufail, F., Leong, H. F., Viola, S., Davis, G. A., Rew, N., Medranda, D., Hofer, M., Simpson, P., Sandrin, M., Chachuat, B., Nelson, J., Renger, T., Murray, J. W., Fantuzzi, A., & Rutherford, A. W. (2025). Locating the missing chlorophylls f in far-red photosystem I. *Science*, eado6830. 10.1126/science.ado6830

Derks, A., Schaven, K., & Bruce, D. (2015). Diverse mechanisms for photoprotection in photosynthesis. Dynamic regulation of photosystem II excitation in response to rapid environmental change. Biochimica et Biophysica Acta (BBA) - Bioenergetics, 1847(4), 468–485. 10.1016/j.bbabio.2015.02.008

Di Valentin, M., & Carbonera, D. (2017). The fine tuning of carotenoid–chlorophyll interactions in light-harvesting complexes: An important requisite to guarantee efficient photoprotection via triplet–triplet energy transfer in the complex balance of the energy transfer processes. *Journal of Physics B: Atomic*, Molecular and Optical Physics, 50(16), 162001. 10.1088/1361-6455/aa7dd4

Domínguez-Martín, M. A., Sauer, P. V., Kirst, H., Sutter, M., Bína, D., Greber, B. J., Nogales, E., Polívka, T., & Kerfeld, C. A. (2022). Structures of a phycobilisome in light-harvesting and photoprotected states. Nature, 609(7928), 835–845. 10.1038/s41586-022-05156-4

Elias, E., Oliver, T. J., & Croce, R. (2024). Oxygenic Photosynthesis in Far-Red Light: Strategies and Mechanisms. Annual Review of Physical Chemistry, 75(1). 10.1146/annurev-physchem-090722-125847

Fiebig, O. C., Harris, D., Wang, D., Hoffmann, M. P., & Schlau-Cohen, G. S. (2023). Ultrafast Dynamics of Photosynthetic Light Harvesting: Strategies for Acclimation Across Organisms. Annual Review of Physical Chemistry, 74(1), 493–520. 10.1146/annurev-physchem-083122-111318

Gan, F., Shen, G., & Bryant, D. A. (2015). Occurrence of far-red light photoacclimation (FaRLiP) in diverse cyanobacteria. Life, 5(1). 10.3390/life5010004

Gan, F., Zhang, S., Rockwell, N. C., Martin, S. S., Lagarias, J. C., & Bryant, D. A. (2014). Extensive remodeling of a cyanobacterial photosynthetic apparatus in far-red light. Science, 345(6202). 10.1126/science.1256963

García-Oneto, T. M., Moyano-Bellido, C., & Domínguez-Martín, M. A. (2024). Structure and function of the light-protective orange carotenoid protein families. Current Research in Structural Biology, 7, 100141. 10.1016/j.crstbi.2024.100141

Genty, B., Briantais, J.-M., & Baker, N. R. (1989). The relationship between the quantum yield of photosynthetic electron transport and quenching of chlorophyll fluorescence. Biochimica et Biophysica Acta (BBA) - General Subjects, 990(1), 87–92. 10.1016/S0304-4165(89)80016-9

Geraldes, V., & Pinto, E. (2021). Mycosporine-Like Amino Acids (MAAs): Biology, Chemistry and Identification Features. Pharmaceuticals, 14(1), 63. 10.3390/ph14010063

Gisriel, C. J. (2024). Recent structural discoveries of photosystems I and II acclimated to absorb far-red light. Biochimica et Biophysica Acta (BBA) - Bioenergetics, 1865(3), 149032. 10.1016/j.bbabio.2024.149032

Gisriel, C. J., Bryant, D. A., Brudvig, G. W., & Cardona, T. (2023). Molecular diversity and evolution of far-red light-acclimated photosystem I. Frontiers in Plant Science, 14. 10.3389/fpls.2023.1289199

Gisriel, C. J., Cardona, T., Bryant, D. A., & Brudvig, G. W. (2022). Molecular Evolution of Far-Red Light-Acclimated Photosystem II. Microorganisms, 10(7), Article 7. 10.3390/microorganisms10071270

Gisriel, C. J., Flesher, D. A., Shen, G., Wang, J., Ho, M. Y., Brudvig, G. W., & Bryant, D. A. (2022b). Structure of a photosystem I-ferredoxin complex from a marine cyanobacterium provides insights into far-red light photoacclimation. Journal of Biological Chemistry, 298(1). 10.1016/j.jbc.2021.101408

Gisriel, C. J., Huang, H.-L., Reiss, K. M., Flesher, D. A., Batista, V. S., Bryant, D. A., Brudvig, G. W., & Wang, J. (2021). Quantitative assessment of chlorophyll types in cryo-EM maps of photosystem I acclimated to far-red light. BBA Advances, 1, 100019. 10.1016/j.bbadva.2021.100019

Gisriel, C. J., Shen, G., Ho, M. Y., Kurashov, V., Flesher, D. A., Wang, J., Armstrong, W. H., Golbeck, J. H., Gunner, M. R., Vinyard, D. J., Debus, R. J., Brudvig, G. W., & Bryant, D. A. (2022a). Structure of a monomeric photosystem II core complex from a cyanobacterium acclimated to far-red light reveals the functions of chlorophylls d and f. Journal of Biological Chemistry, 298(1). 10.1016/j.jbc.2021.101424

Gisriel, C., Shen, G., Kurashov, V., Ho, M.-Y., Zhang, S., Williams, D., Golbeck, J. H., Fromme, P., & Bryant, D. A. (2020). The structure of Photosystem I acclimated to far-red light illuminates an ecologically important acclimation process in photosynthesis. Science Advances, 6(6), eaay6415. 10.1126/sciadv.aay6415

Gobets, B., & Van Grondelle, R. (2001). Energy transfer and trapping in photosystem I. Biochimica et Biophysica Acta (BBA) - Bioenergetics, 1507(1–3), 80–99. 10.1016/S0005-2728(01)00203-1

Hashimoto, H., Uragami, C., & Cogdell, R. J. (2016). Carotenoids and Photosynthesis. In C. Stange (Ed.), Carotenoids in Nature (Vol. 79, pp. 111–139). Springer International Publishing. 10.1007/978-3-319-39126-7_4

Ho, J., Kish, E., Méndez-Hernández, D. D., WongCarter, K., Pillai, S., Kodis, G., Niklas, J., Poluektov, O. G., Gust, D., Moore, T. A., Moore, A. L., Batista, V. S., & Robert, B. (2017). Triplet–triplet energy transfer in artificial and natural photosynthetic antennas. Proceedings of the National Academy of Sciences, 114(28). 10.1073/pnas.1614857114

Ho, M. Y., Gan, F., Shen, G., & Bryant, D. A. (2017). Far-red light photoacclimation (FaRLiP) in Synechococcus sp. PCC 7335. II.Characterization of phycobiliproteins produced during acclimation to far-red light. Photosynthesis Research, 131(2). 10.1007/s11120-016-0303-5

Karapetyan, N. V., Schlodder, E., Van Grondelle, R., & Dekker, J. P. (2006). The Long Wavelength Chlorophylls of Photosystem I. In J. H. Golbeck (Ed.), Photosystem I (Vol. 24, pp. 177–192). Springer Netherlands. 10.1007/978-1-4020-4256-0_13

Kato, K., Shinoda, T., Nagao, R., Akimoto, S., Suzuki, T., Dohmae, N., Chen, M., Allakhverdiev, S. I., Shen, J.-R., Akita, F., Miyazaki, N., & Tomo, T. (2020). Structural basis for the adaptation and function of chlorophyll f in photosystem I. Nature Communications, 11(1), 238. 10.1038/s41467-019-13898-5

Kerfeld, C. A. (2004). Structure and Function of the Water-Soluble Carotenoid-Binding Proteins of Cyanobacteria. Photosynthesis Research, 81(3), 215–225. 10.1023/B:PRES.0000036886.60187.c8

Kramer, D. M., Johnson, G., Kiirats, O., & Edwards, G. E. (2004). New Fluorescence Parameters for the Determination of Q_A_ Redox State and Excitation Energy Fluxes. Photosynthesis Research, 79(2), 209–218. 10.1023/B:PRES.0000015391.99477.0d

Latifi, A., Ruiz, M., & Zhang, C.-C. (2009). Oxidative stress in cyanobacteria. FEMS Microbiology Reviews, 33(2), 258–278. 10.1111/j.1574-6976.2008.00134.x

Li, Y., Scales, N., Blankenship, R. E., Willows, R. D., & Chen, M. (2012). Extinction coefficient for red-shifted chlorophylls: Chlorophyll d and chlorophyll f. Biochimica et Biophysica Acta (BBA) - Bioenergetics, 1817(8), 1292–1298. 10.1016/j.bbabio.2012.02.026

Liistro, E., Battistuzzi, M., Storti, M., Boccia, B., Cocola, L., Perin, G., Morosinotto, T., & La Rocca, N. (2025). Thylakoids reorganization enables driving photosynthesis under far-red light in the microalga *Nannochloropsis gaditana*. New Phytologist. 10.1111/nph.70786

Llewellyn, C. A., Greig, C., Silkina, A., Kultschar, B., Hitchings, M. D., & Farnham, G. (2020). Mycosporine-like amino acid and aromatic amino acid transcriptome response to UV and far-red light in the cyanobacterium *Chlorogloeopsis fritschii* PCC 6912. Scientific Reports, 10(1), 20638. 10.1038/s41598-020-77402-6

Mascoli, V., Bersanini, L., & Croce, R. (2020). Far-red absorption and light-use efficiency trade-offs in chlorophyll f photosynthesis. Nature Plants, 6(8). 10.1038/s41477-020-0718-z

Mascoli, V., Bhatti, A. F., Bersanini, L., van Amerongen, H., & Croce, R. (2022). The antenna of far-red absorbing cyanobacteria increases both absorption and quantum efficiency of Photosystem II. Nature Communications, 13(1). 10.1038/s41467-022-31099-5

Melis, A. (1999). Photosystem-II damage and repair cycle in chloroplasts: What modulates the rate of photodamage *in vivo*? Trends in Plant Science, 4(4), 130–135. 10.1016/S1360-1385(99)01387-4

Miyashita, H., Ikemoto, H., Kurano, N., Adachi, K., Chihara, M., & Miyachi, S. (1996). Chlorophyll d as a major pigment. Nature, 383(6599), 402–402. 10.1038/383402a0

Moran, R. (1982). Formulae for Determination of Chlorophyllous Pigments Extracted with N,N - Dimethylformamide. Plant Physiology, 69(6). 10.1104/pp.69.6.1376

Morosinotto, T., Breton, J., Bassi, R., & Croce, R. (2003). The Nature of a Chlorophyll Ligand in Lhca Proteins Determines the Far Red Fluorescence Emission Typical of Photosystem I. Journal of Biological Chemistry, 278(49), 49223–49229. 10.1074/jbc.M309203200

Muzzopappa, F., & Kirilovsky, D. (2020). Changing Color for Photoprotection: The Orange Carotenoid Protein. Trends in Plant Science, 25(1), 92–104. 10.1016/j.tplants.2019.09.013

Muzzopappa, F., Wilson, A., & Kirilovsky, D. (2019). Interdomain interactions reveal the molecular evolution of the orange carotenoid protein. Nature Plants, 5(10), 1076–1086. 10.1038/s41477-019-0514-9

Niziński, S., Hartmann, E., Shoeman, R. L., Tarnawski, M., Wilson, A., Reinstein, J., Kirilovsky, D., Sliwa, M., Burdziński, G., & Schlichting, I. (2025). Two-Photon-Driven Photoprotection Mechanism in Echinenone-Functionalized Orange Carotenoid Protein. Journal of the American Chemical Society, 147(5), 4100–4110. 10.1021/jacs.4c13341

Nürnberg, D. J., Morton, J., Santabarbara, S., Telfer, A., Joliot, P., Antonaru, L. A., Ruban, A. V., Cardona, T., Krausz, E., Boussac, A., Fantuzzi, A., & Rutherford, A. W. (2018). Photochemistry beyond the red limit in chlorophyll f–containing photosystems. Science, 360(6394), 1210–1213. 10.1126/science.aar8313

Polívka, T., & Sundström, V. (2004). Ultrafast Dynamics of Carotenoid Excited States−From Solution to Natural and Artificial Systems. Chemical Reviews, 104(4), 2021–2072. 10.1021/cr020674n

Punginelli, C., Wilson, A., Routaboul, J.-M., & Kirilovsky, D. (2009). Influence of zeaxanthin and echinenone binding on the activity of the Orange Carotenoid Protein. Biochimica et Biophysica Acta (BBA) - Bioenergetics, 1787(4), 280–288. 10.1016/j.bbabio.2009.01.011

Rajneesh, ., Pathak, J., Chatterjee, A., Singh, S., & Sinha, R. (2017). Detection of Reactive Oxygen Species (ROS) in Cyanobacteria Using the Oxidant-sensing Probe 2’,7’-Dichlorodihydrofluorescein Diacetate (DCFH-DA). BIO-PROTOCOL, 7(17). 10.21769/BioProtoc.2545

Ramel, F., Birtic, S., Cuiné, S., Triantaphylidès, C., Ravanat, J.-L., & Havaux, M. (2012). Chemical Quenching of Singlet Oxygen by Carotenoids in Plants1[C][W]. Plant Physiology, 158(3), 1267–1278. 10.1104/pp.111.182394

Rippka, R., Deruelles, J., & Waterbury, J. B. (1979). Generic assignments, strain histories and properties of pure cultures of cyanobacteria. Journal of General Microbiology, 111(1). 10.1099/00221287-111-1-1

Santabarbara, S., Agostini, G., Casazza, A. P., Syme, C. D., Heathcote, P., Böhles, F., Evans, M. C. W., Jennings, R. C., & Carbonera, D. (2007). Chlorophyll triplet states associated with Photosystem I and Photosystem II in thylakoids of the green alga *Chlamydomonas reinhardtii*. Biochimica et Biophysica Acta (BBA) - Bioenergetics, 1767(1), 88–105. 10.1016/j.bbabio.2006.10.007

Schmitt, F.-J., & Friedrich, T. (2024). Adaptation processes in Halomicronema hongdechloris, an example of the light-induced optimization of the photosynthetic apparatus on hierarchical time scales. Frontiers in Plant Science, 15, 1359195. 10.3389/fpls.2024.1359195

Sedoud, A., López-Igual, R., Ur Rehman, A., Wilson, A., Perreau, F., Boulay, C., Vass, I., Krieger-Liszkay, A., & Kirilovsky, D. (2014). The Cyanobacterial Photoactive Orange Carotenoid Protein Is an Excellent Singlet Oxygen Quencher. The Plant Cell, 26(4), 1781–1791. 10.1105/tpc.114.123802

Shen, L., Zhang, Z., Zhang, L., Huang, D., Yu, G., Chen, M., Li, R., & Qiu, B. (2025). Widespread distribution of chlorophyll *f* [producing *Leptodesmis* cyanobacteria. Journal of Phycology, 61(1), 144–160. 10.1111/jpy.13538

Silkina, A., Kultschar, B., & Llewellyn, C. A. (2019). Far-Red Light Acclimation for Improved Mass Cultivation of Cyanobacteria. Metabolites, 9(8), 170. 10.3390/metabo9080170

Singh, S. P., Rastogi, R. P., Häder, D.-P., & Sinha, R. P. (2014). Temporal dynamics of ROS biogenesis under simulated solar radiation in the cyanobacterium *Anabaena variabilis* PCC 7937. Protoplasma, 251(5), 1223–1230. 10.1007/s00709-014-0630-3

Slonimskiy, Y. B., Zupnik, A. O., Varfolomeeva, L. A., Boyko, K. M., Maksimov, E. G., & Sluchanko, N. N. (2022). A primordial Orange Carotenoid Protein: Structure, photoswitching activity and evolutionary aspects. International Journal of Biological Macromolecules, 222, 167–180. 10.1016/j.ijbiomac.2022.09.131

Soulier, N., Laremore, T. N., & Bryant, D. A. (2020). Characterization of cyanobacterial allophycocyanins absorbing far-red light. Photosynthesis Research, 145(3), 189–207. 10.1007/s11120-020-00775-2

Trinugroho, J. P., Bečková, M., Shao, S., Yu, J., Zhao, Z., Murray, J. W., Sobotka, R., Komenda, J., & Nixon, P. J. (2020). Chlorophyll f synthesis by a super-rogue photosystem II complex. Nature Plants, 6(3). 10.1038/s41477-020-0616-4

Waditee-Sirisattha, R., & Kageyama, H. (2022). Extremophilic cyanobacteria. In Cyanobacterial Physiology: From Fundamentals to Biotechnology. 10.1016/B978-0-323-96106-6.00012-5

Wilson, A., Ajlani, G., Verbavatz, J.-M., Vass, I., Kerfeld, C. A., & Kirilovsky, D. (2006). A Soluble Carotenoid Protein Involved in Phycobilisome-Related Energy Dissipation in Cyanobacteria. The Plant Cell, 18(4), 992–1007. 10.1105/tpc.105.040121

Wilson, A., Gwizdala, M., Mezzetti, A., Alexandre, M., Kerfeld, C. A., & Kirilovsky, D. (2012). The Essential Role of the N-Terminal Domain of the Orange Carotenoid Protein in Cyanobacterial Photoprotection: Importance of a Positive Charge for Phycobilisome Binding. The Plant Cell, 24(5), 1972–1983. 10.1105/tpc.112.096909

Wolf, B. M., & Blankenship, R. E. (2019). Far-red light acclimation in diverse oxygenic photosynthetic organisms. Photosynthesis Research, 142(3), 349–359. 10.1007/s11120-019-00653-6

Zampieri, R. M., Bizzotto, E., Campanaro, S., Caldara, F., Bellucci, M., & La Rocca, N. (2025). *Kovacikia euganea* sp. Nov. (Leptolyngbyaceae, Cyanobacteria), a new chlorophyll f producing cyanobacterium from the Euganean Thermal District (Italy). Frontiers in Microbiology, 16. 10.3389/fmicb.2025.1545008

