## Supplementary Material for "Enhanced carotenoid photoprotection in Far-Red light acclimated *Chroococcidiopsis thermalis*"

a

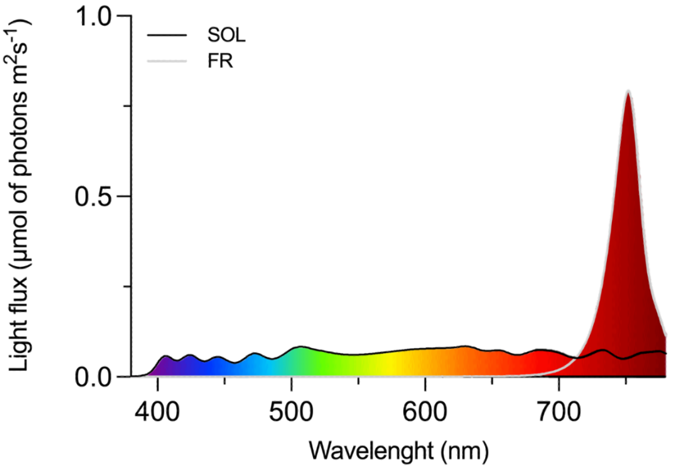


b

| Wavebands | SOL | FR |
|---|---|---|
| **UV light (380-400 nm)** µmol of photons m$^2$s$^{-1}$ | 0.18 | 0.01 |
| **Blue light (400-500 nm)** µmol of photons m$^2$s$^{-1}$ | 5.29 | 0.05 |
| **Green light (500-600 nm)** µmol of photons m$^2$s$^{-1}$ | 7.07 | 0.05 |
| **Red light (600-700 nm)** µmol of photons m$^2$s$^{-1}$ | 7.45 | 0.29 |
| **Far red light (700-780 nm)** µmol of photons m$^2$s$^{-1}$ | 5.01 | 24.60 |
| **Total VIS light (380-700 nm)** µmol of photons m$^2$s$^{-1}$ | 19.99 | 0.41 |
| **Total light (380-780 nm)** µmol of photons m$^2$s$^{-1}$ | 25 | 25 |

Figure S1. Irradiance conditions used in this study. a) solar (SOL) and far red (FR) light spectra to which PCC7203 cells were exposed, in terms of µmol of photons m$^2$s$^{-1}$, b) Distribution of light in the two spectra used in this work: SOL (solar-like) and FR (far red light). In the table is indicated the amount of µmol of photons m$^2$s$^{-1}$ provided in the wavebands UV (380-400 nm), blue (400-500 nm), green (500-600 nm), red (600-700 nm), far red (700-780 nm); the total amount of visible (380-700, VIS) light and total (380-780 nm) light reported. SOL: solar; FR: far red; UV: ultraviolet; VIS: visible.

### Principles of ODMR technique

Optically Detected Magnetic Resonance spectroscopy is particularly suited to investigate photogenerated triplet states, as it combines the high *selectivity* obtained through microwave-induced transitions between spin sublevels with the high *sensitivity* of optical detection.

Triplet states are molecular species characterised by two unpaired electrons in their electronic molecular configuration. As the name suggests, the interactions (dipolar couplings) between the unpaired electrons make the triplet state split in three different sublevels, which are not degenerate, even without the application of an external perturbation such as a static magnetic field. Triplet sublevels are commonly called $T_X$, $T_Y$ and $T_Z$. Their energies are dependent on the properties of the spin dipolar couplings within the triplet state, and can be described by the zero-filed splitting (ZFS) parameters |D| and |E|.

Considering photosynthetic systems, chlorophyll triplet states can be photogenerated by two main mechanisms, either via Intersystem Crossing (ISC, typically for antenna chlorophylls) or through recombination processes (chlorophylls participating in electron transfer inside PSI and PSII reaction centres).

Carotenoids can populate triplet states too, but only through a mechanism called Triplet-Triplet Energy Transfer (TTET). Briefly, if the geometrical conditions of sufficient proximity and correct orientation are fulfilled, a carotenoid can populate its triplet state by inheriting it from an adjacent chlorophyll that is successfully coupled to it. In this sense, chlorophylls can act as triplet donors, while carotenoids act as triplet acceptors. This mechanism is at the basis of the photoprotection activity exerted by carotenoids. In fact, chlorophyll triplets can photosensitize ROS, while carotenoid triplets cannot. If the chlorophyll-to-carotenoid triplet transfer is fast enough, energy can be safely dissipated before becoming dangerous, especially in the conditions of luminous overexposure leading to a saturated photosynthetic turnover. Detection of carotenoid triplets in photosynthetic systems is thus a fingerprint of their photoprotective action.

To understand the working principles of the ODMR technique, with a focus on the mechanism that makes it possible to detect carotenoid triplets by monitoring variations in the chlorophyll fluorescence, Figure S2 and S3 are reported below. In the condition of continuous illumination, chlorophylls populate triplet sublevels with (anisotropic) populating rate constants assumed as $k_Y > k_X > k_Z$. The green circles represent chlorophyll triplet sublevel populations. Through TTET, carotenoids populate their triplet state, with sublevel populations indicated by the orange circles. In final, carotenoids relax to ground state with (anisotropic) depopulation rate constants assumed as $p_Y > p_Z > p_X$. This situation represents the case in which no microwaves are sent to the sample (Figure S2).

When high-power microwave frequencies are in resonance with a carotenoid triplet sublevel transition, for example the 2|E| transition indicated with a purple arrow in Figure S3, the triplet energy levels that are put in resonance can equalize their populations. For this example, carotenoid $T_Y$ increases its population, while $T_X$ decreases it.

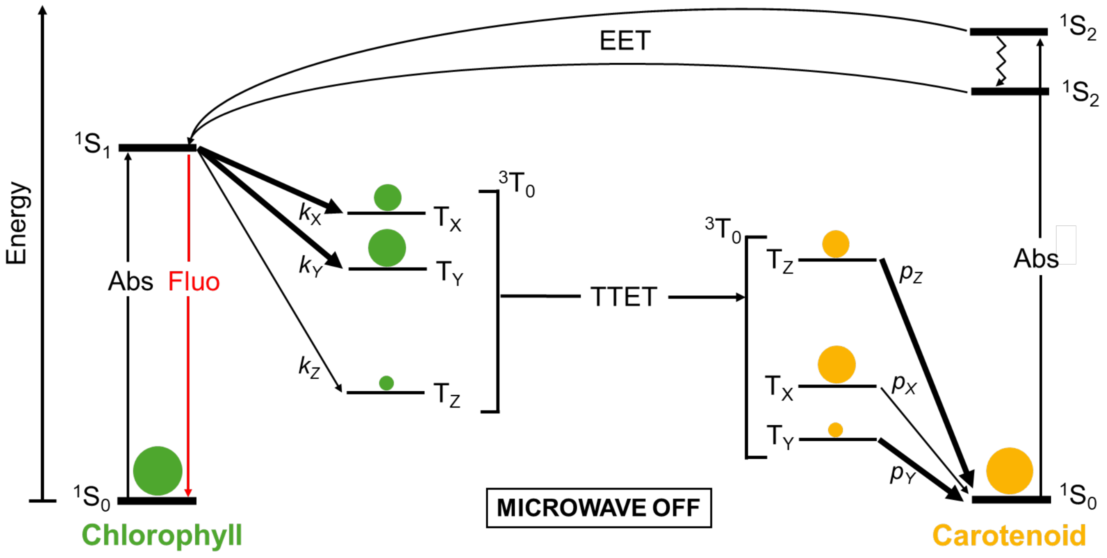


**Figure S2**. Jablonski diagram for the coupled chlorophyll-carotenoid system without application of microwaves.

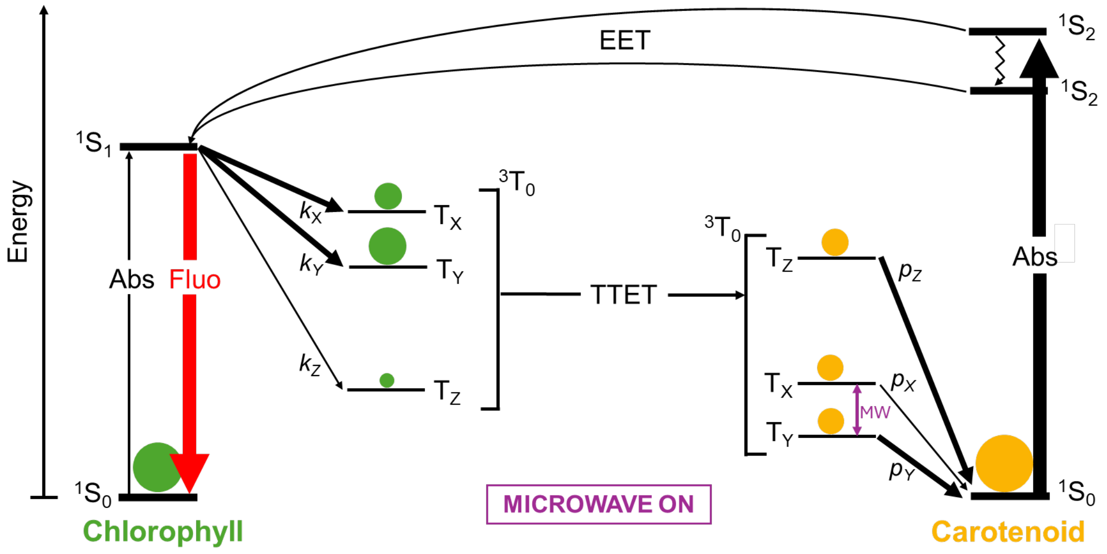


**Figure S3.** Jablonski diagram for the coupled chlorophyll-carotenoid system with the application of resonant microwaves promoting the 2|E| carotenoid triplet transition.

Considering that $p_Y > p_Z$, the larger population of $T_Y$ induced by the microwaves action will also have the effect of augmenting the population of the carotenoid ground state $^1S_0$. In a “cascade effect”, a larger ground state carotenoid population will induce enhanced carotenoid absorption (indicated by the thickened black arrow in Figure S3) and, as a consequence of excitation energy transfer (EET), enhanced chlorophyll fluorescence (thickened red arrow in Figure S3). This explains why carotenoid triplet transitions can affect chlorophyll fluorescence, if the two pigments are coupled.

Combining optical detection and phase sensitive (Lock-In) amplification of the microwave on/off modulation, high sensitivity can be reached in this type of ODMR experiment. Most importantly, by monitoring the magnetic properties (microwave frequency pumping) and optical properties (optical wavelength range of detection) simultaneously, a correlation between these two aspects can be assessed for the studied systems, giving useful information on their structural and energetic characteristics.

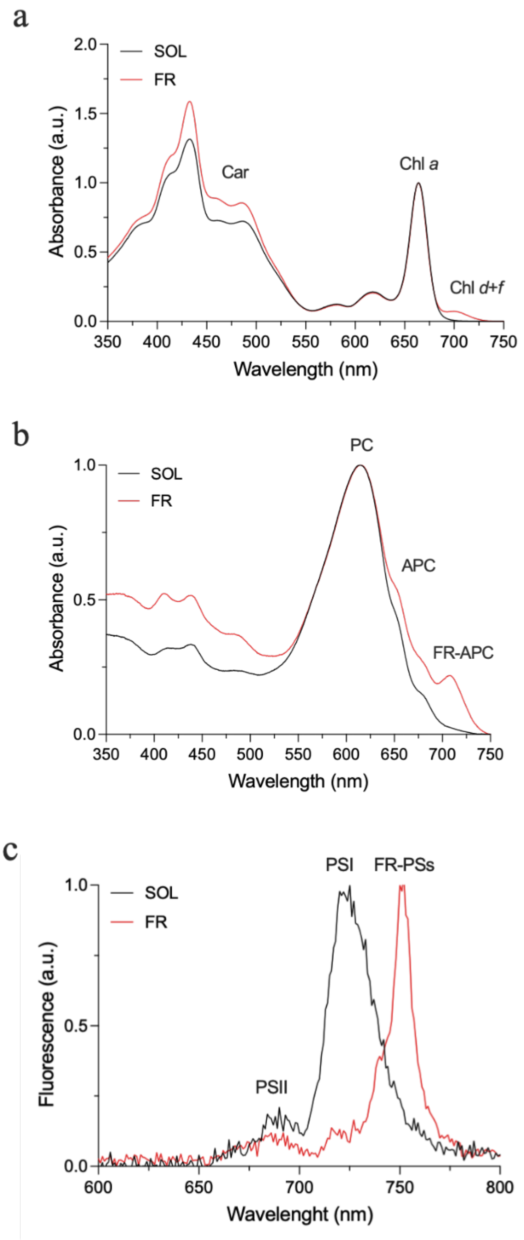


**Figure S4**. Pigment composition and low temperature (77 K) fluorescence emission of *C. thermalis* cells acclimated to SOL and FR light after 21 days of exposure. a) absorption spectra of chlorophylls and carotenoids extracts normalized at 660 nm; b) absorption spectra of phycobiliproteins extracts normalized at 620 nm c) low temperature (77 K) fluorescence emission spectra of cells excited at 440 nm, normalized to maximum emission peak of each sample. Reported spetra are the mean of three biological replicates. SOL: Solar light; FR: Far-red light; Car: Total carotenoids; Chl *a*: Chlorophyll *a*; Chl *d*+*f*: Chlorophyll *d* and *f*; PC: Phycocyanin; APC: Allophycocyanin; FR-APC: Far-Red Allophycocyanin; PSII: Photosystem II; PSI: Photosystem I; FR-PS: Far-red-acclimated photosystems I and II.

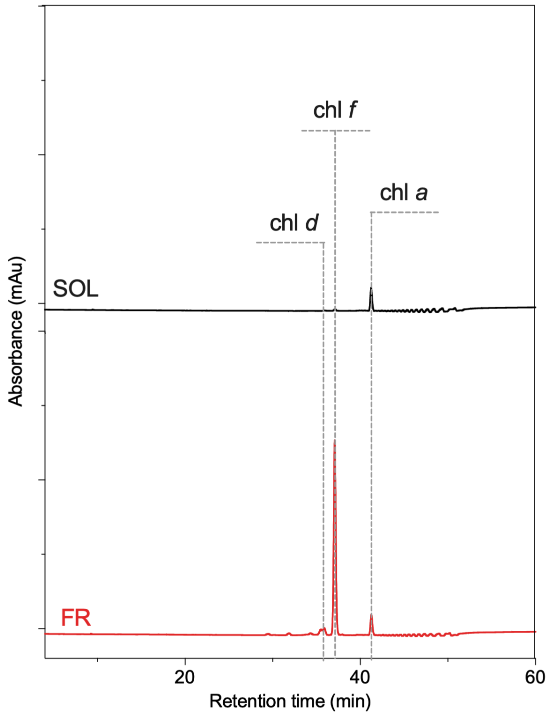


**Figure S5**. Chromatogram at 705 nm of extracts from cells acclimated to SOL (black) and FR (red) light. Dotted lines indicate the retention times of the detected pigments. Chl *d*: chlorophyll *d*; chl *f*: chlorophyll *f*; chl *a*: chlorophyll *a*; SOL: solar; FR: far red.

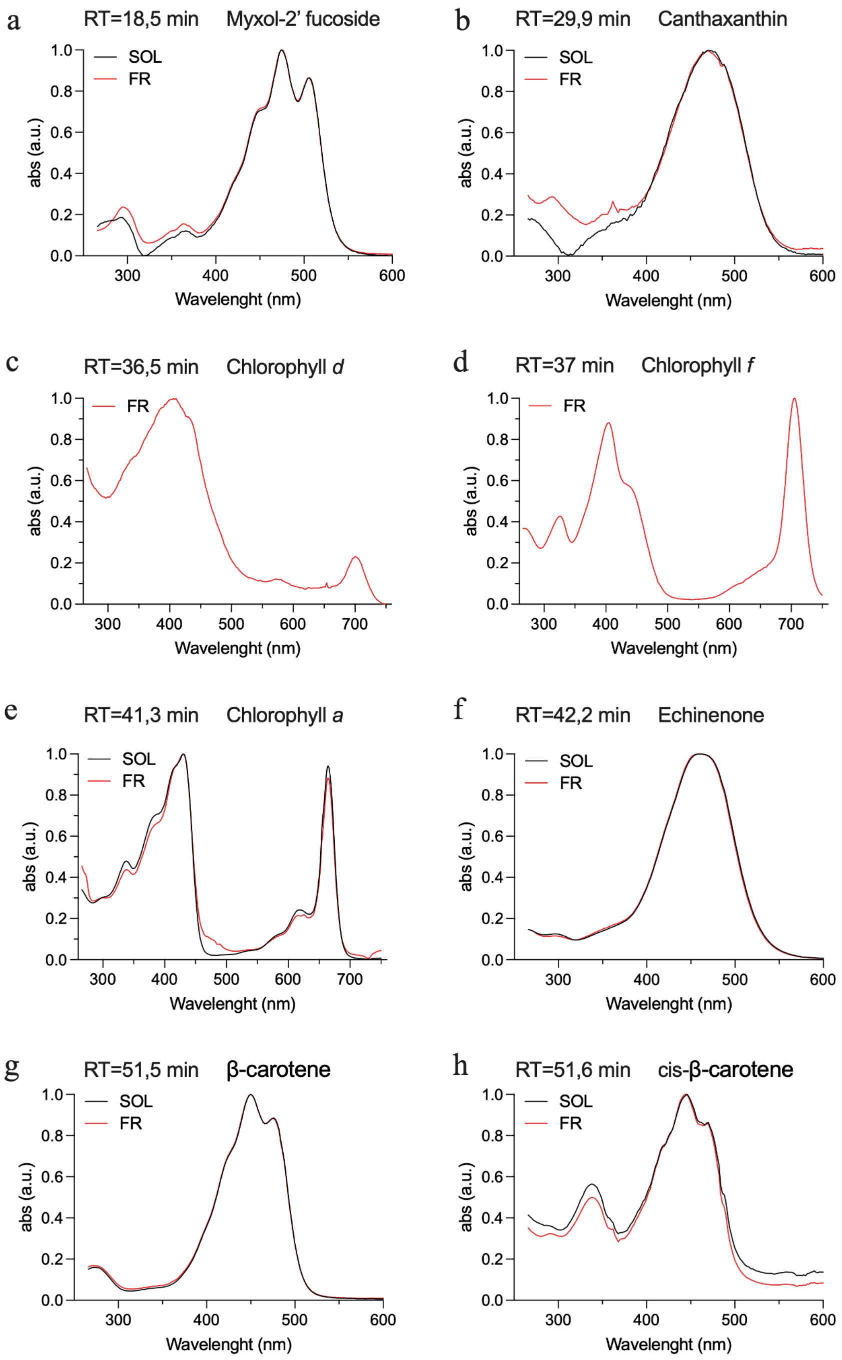


**Figure S6**. Spectra of pigments corresponding to HPLC chromatogram peaks, detected with the DAD. a) Mixoxanthophyll, eluted at minute 18,5; b) cantaxanthin, eluted at minute 29,9; chlorophyll *d*, eluted at minute 36,5 only in extracts of FR acclimated cells; d) chlorophyll *f*, eluted at minute 37 only in extracts of FR acclimated cells; e) chlorophyll *a*, eluted at minute 41,3, normalized at 664 nm; f) echinenone, eluted at minute 42,2; g) β-carotene, eluted at minute 51,5; cis β-carotene, eluted at minute 51,6. SOL: solar, FR: far-red; RT: retention time.

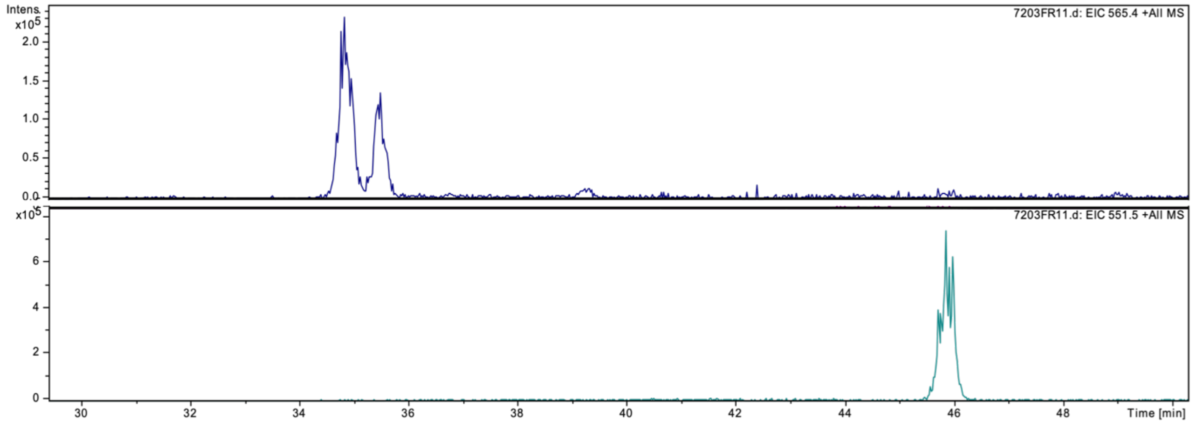


**Figure S7**. Extracted ion currents for m/z ions compatible with canthaxanthin and echinenone. The mass spectrometry analysis reports the extracted ion current generated from molecular species having m/z ratios of 565.4 (top purple trace) and 551.5 (green bottom trace), respectively assigned to two isomers of canthaxanthin and echinenone.
